# Multiple independent acquisitions of ACE2 usage in MERS-related coronaviruses

**DOI:** 10.1101/2023.10.02.560486

**Authors:** Cheng-Bao Ma, Chen Liu, Young-Jun Park, Jingjing Tang, Jing Chen, Qing Xiong, Jimin Lee, Cameron Stewart, Daniel Asarnow, Jack Brown, M. Alejandra Tortorici, Xiao Yang, Ye-Hui Sun, Yuan-Mei Chen, Xiao Yu, Jun-Yu Si, Peng Liu, Fei Tong, Mei-Ling Huang, Jing Li, Zheng-Li Shi, Zengqin Deng, David Veesler, Huan Yan

**Affiliations:** State Key Laboratory of Virology, College of Life Sciences, TaiKang Center for Life and Medical Sciences, Wuhan University; Wuhan, Hubei, 430072, China; Department of Biochemistry, University of Washington; Seattle, WA 98195, USA; Howard Hughes Medical Institute, University of Washington; Seattle, WA 98195, USA; Key Laboratory of Virology and Biosafety, Wuhan Institute of Virology, Chinese Academy of Sciences; Wuhan, China; Guangzhou Laboratory, Guangzhou International Bio Island; Guangzhou, China; Hubei Jiangxia Laboratory; Wuhan, China

## Abstract

The angiotensin-converting enzyme 2 (ACE2) receptor is shared by various coronaviruses with distinct receptor-binding domain (RBD) architectures, yet our understanding of these convergent acquisition events remains elusive. Here, we report that two European bat MERS-related coronaviruses (MERSr-CoVs) infecting *Pipistrellus nathusii* (P.nat), MOW15-22 and PnNL2018B, use ACE2 as their receptor, with narrow ortholog specificity. Cryo-electron microscopy structures of the MOW15-22 RBD-ACE2 complex unveil an unexpected and entirely distinct binding mode, mapping 50Å away from that of any other known ACE2-using coronaviruses. Functional profiling of ACE2 orthologs from 105 mammalian species led to the identification of host tropism determinants, including an ACE2 N432-glycosylation restricting viral recognition, and the design of a soluble P.nat ACE2 mutant with potent viral neutralizing activity. Our findings reveal convergent acquisition of ACE2 usage for merbecoviruses found in European bats, underscoring the extraordinary diversity of ACE2 recognition modes among coronaviruses and the promiscuity of this receptor.

**Graphic abstract:** 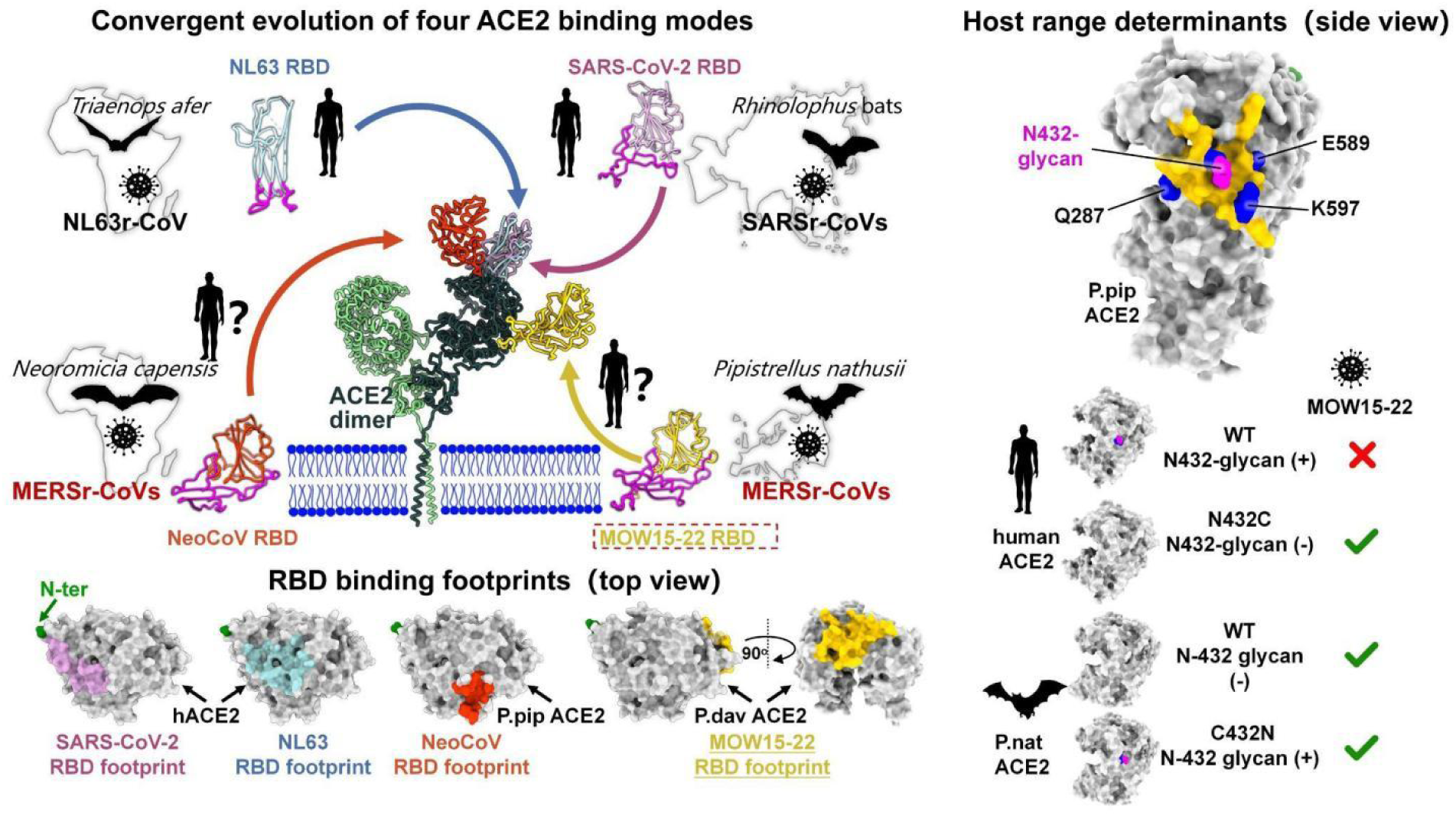

## INTRODUCTION

Human MERS-CoV belongs to the *Merbecovirus* subgenus (also known as lineage C β- coronavirus or group 2c coronavirus) and is the causative agent of Middle-East respiratory syndrome (MERS) with a case-fatality rate of 36%.^1^ The evolutionary trajectory and host- switching history of MERS-CoV remain unclear. While dromedary camels are established intermediate hosts of MERS-CoV, increasing evidence from newly identified viral sequences indicates that Old World vesper bats (Vespertilionidae) are natural merbecovirus reservoirs.^2–7^ However, no currently known bat MERS-related coronaviruses (MERSr-CoVs) exhibit close similarity to human or camel MERS-CoV at the whole-genome level.^8,9^ NeoCoV, identified in African Laephotis bats (Cape serotine), represents the closest known relative to MERS-CoV, sharing 85.5% whole genome nucleotide sequence identity.^10^

Receptor usage determines host tropism and transmission of coronaviruses^11^. Dipeptidyl peptidase-4 (DPP4) has been documented as the entry receptor for several merbecoviruses, including MERS-CoV, HKU4, MjHKU4r, HKU25, and BtCoV-422.^12–18^ NeoCoV and the closely related PDF-2180 are two bat MERSr-CoVs that harbor markedly different spike (S) glycoproteins relative to that of MERS-CoV, and their cognate receptor remained elusive for a decade. For a long time, ACE2 was solely considered the host receptor recognized by members of sarbecoviruses (e.g., SARS-CoV-1 and SARS-CoV-2) and setracoviruses (e.g., NL63) through two different ACE2-binding modes with largely overlapping footprints in spite of their distinct receptor-binding domain (RBD) architectures.^19–23^ We recently revealed that NeoCoV and PDF-2180 also use the ACE2 receptor with a broad tropism across mammals but low efficiency in using human ACE2 (hACE2).^9,24^ Cryo-electron microscopy (cryo-EM) analysis of the NeoCoV and the PDF-2180 RBDs in complex with *Pipistrellus pipistrellus* (P.pip) ACE2 unveiled a distinct binding mode from that of SARS-CoV-1/SARS-CoV-2 and of NL63 involving extensive interactions with ACE2 N linked-glycans. The footprint of their RBDs shared few residues with that of SARS-CoV-1/2 or NL63, highlighting a convergent evolutionary history of ACE2 acquisition in these viruses.^9^ This discovery suggested that a receptor switch might have occurred during MERS-CoV emergence in animals, such as bats and camels, potentially through recombination between a NeoCoV-like MERSr-CoV and a DPP4- using merbecovirus (e.g., HKU4 or BtCoV-422).^8,13^ Furthermore, these findings underscored the complexity of receptor binding modes used by merbecoviruses with diverse RBD sequences, many of which remain enigmatic.

Here, we discovered that two MERSr-CoVs circulating in European bats, designated MOW15-22 and PnNL2018B, utilize a subset of mammalian ACE2 orthologs as entry receptors. Cryo-EM analyses reveal that these viruses recognize ACE2 through an unprecedented binding mode, mapping 50Å away from the footprint of any other coronaviruses, emphasizing multiple independent acquisitions of ACE2 usage in bat MERSr-CoVs. This study sheds light on the global diversity and distribution of ACE2-using merbecoviruses and unexpected convergent evolution events, underscoring the zoonotic potential associated with these viruses.

## RESULTS

### Prediction of a distinct receptor recognition mode utilized by two MERSr-CoVs

In light of the unexpected ACE2 usage of NeoCoV and PDF-2180, we investigated whether other merbecoviruses utilize ACE2 as a receptor. We conducted phylogenetic analyses of representative merbecovirus sequences, with a special emphasis on viruses belonging to MERSr- CoVs, as defined by amino acid sequence identity of the five concatenated replicase domains (3CLpro, NiRAN, RdRp, ZBD, and HEL1) greater than 92.4% compared with MERS-CoV (Figure 1A).^25^ At the whole-genome level, MOW15-22 and PnNL2018B (formerly PN-βCoV) are two bat MERSr-CoVs with the highest genetic similarity to MERS-CoV after NeoCoV and PDF-2180 (Figure 1A). At the S glycoprotein level, the MOW15-22 and PnNL2018B S glycoproteins cluster together and are distantly related to all other MERSr-CoVs. By contrast, hedgehog merbecovirus (Erinaceus coronavirus, EriCoV) HKU31 exhibits the highest sequence identity with NeoCoV and PDF-2180, although HKU31 is not known to use ACE2 (Figure 1B).^9^ Simplot analysis comparing several viral genome sequences with MOW15-22 reveals both MERS-CoV and NeoCoV display an extensive divergence within the S1 subunit region (Figure 1C). Consistent with the phylogenetic tree based on S glycoproteins, pairwise analysis shows MOW15-22 and PnNL2018B share only 31-36% and 57-60% amino acid sequence identities with RBDs and S glycoproteins of other merbecoviruses, respectively (Figure 1D). Alignment of the region corresponding to the NeoCoV receptor-binding motif (RBM) reveals the presence of two insertions and a disulfide bond specific of MOW15-22 and PnNL2018B (Figure 1E). Only two (positions N502MOW15-22 and G566MOW15-22) out of nine NeoCoV residues critical for interactions with P.pip ACE2 are conserved with MOW15-22 and PnNL2018B (Figure 1E).^9^ Furthermore, the two insertions and elongated α-helix in the RBM present in AlphaFold2 predicted structures^26^ potentially affect receptor recognition (Figure 1F). Overall, these distinct sequence and structural features indicate that MOW15-22 and PnNL2018B may utilize a previously uncharacterized receptor binding mode.

**Figure 1.**
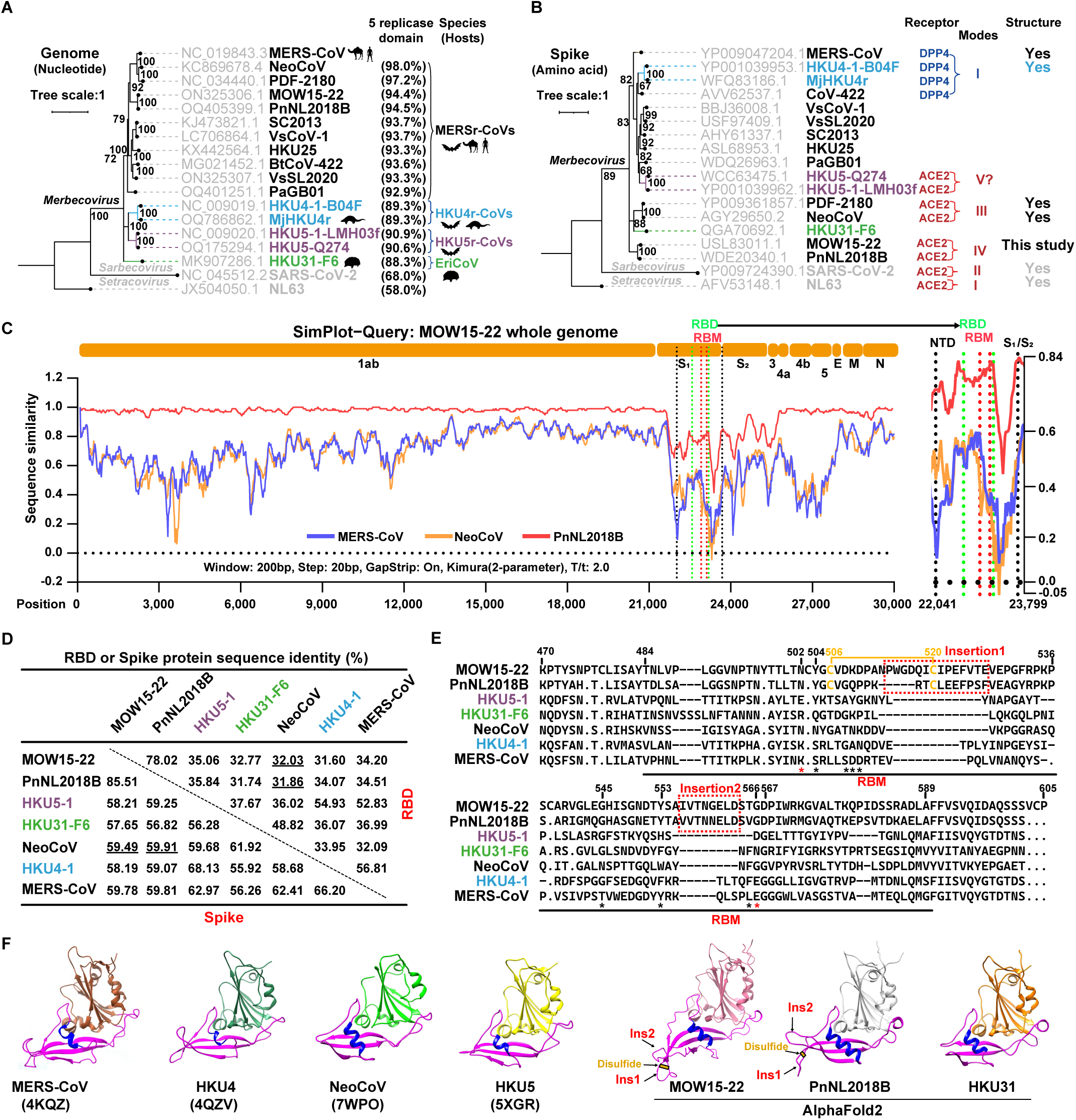
Prediction of a distinct receptor recognition mode utilized by two MERSr-CoVs. (**A-B**) Phylogenetic trees of representative merbecoviruses were generated using complete genome nucleotide sequences (A) or S glycoprotein amino acid sequences (B) with the IQ-tree method. NL63 was set as an outgroup. Amino acid sequence identities of five replicase domains for coronavirus classification, host species, and receptor-related information (receptor usage, binding mode, and availability of RBD/receptor complex structure) are indicated. (**C**) Simplot analysis of the complete genome sequence similarity of several MERSr-CoVs analyzed based on the MOW15-22 genome. The right panel magnifies the RBD and adjacent regions. (**D**) Pairwise RBD and S amino acid sequence identities of indicated merbecoviruses. (**E**) RBM sequence alignment of the indicated merbecoviruses. MOW15-22 and PnNL2018B-specific insertions and disulfides are indicated in red and yellow, respectively. Red/black asterisks: residues crucial for NeoCoV interactions with P.pip ACE2 that are conserved/not conserved with MOW15-22 and PnNL2018B. The MOW15-22 residue numbering is shown. (**F**) Cryo-EM structures or AlphaFold2-predicted RBD structures of representative merbecoviruses. Magenta indicates putative RBMs, blue represents the RBM helix, and the MOW15-22 and PnNL2018B-specific insertions (ins) are indicated. The yellow sticks represent specific disulfide bonds in RBM insertion 1.

### MOW15-22 and PnNL2018B use ACE2 as receptor

MOW15-22 and PnNL2018B were recently sampled in *Pipistrellus nathusii* (P.nat, the common host) in Russia (Moscow region) and the Netherlands, respectively (Figure 2A).^27,28^ P.nat inhabits a wide range of Europe and undertakes seasonal long-distance migrations, typically from northeast to southwest Europe.^29–31^ Previous reports have proposed that DPP4 may serve as the entry receptor for MOW15-22 and PnNL2018B based on molecular docking analyses.^27,28^

**Figure 2.**
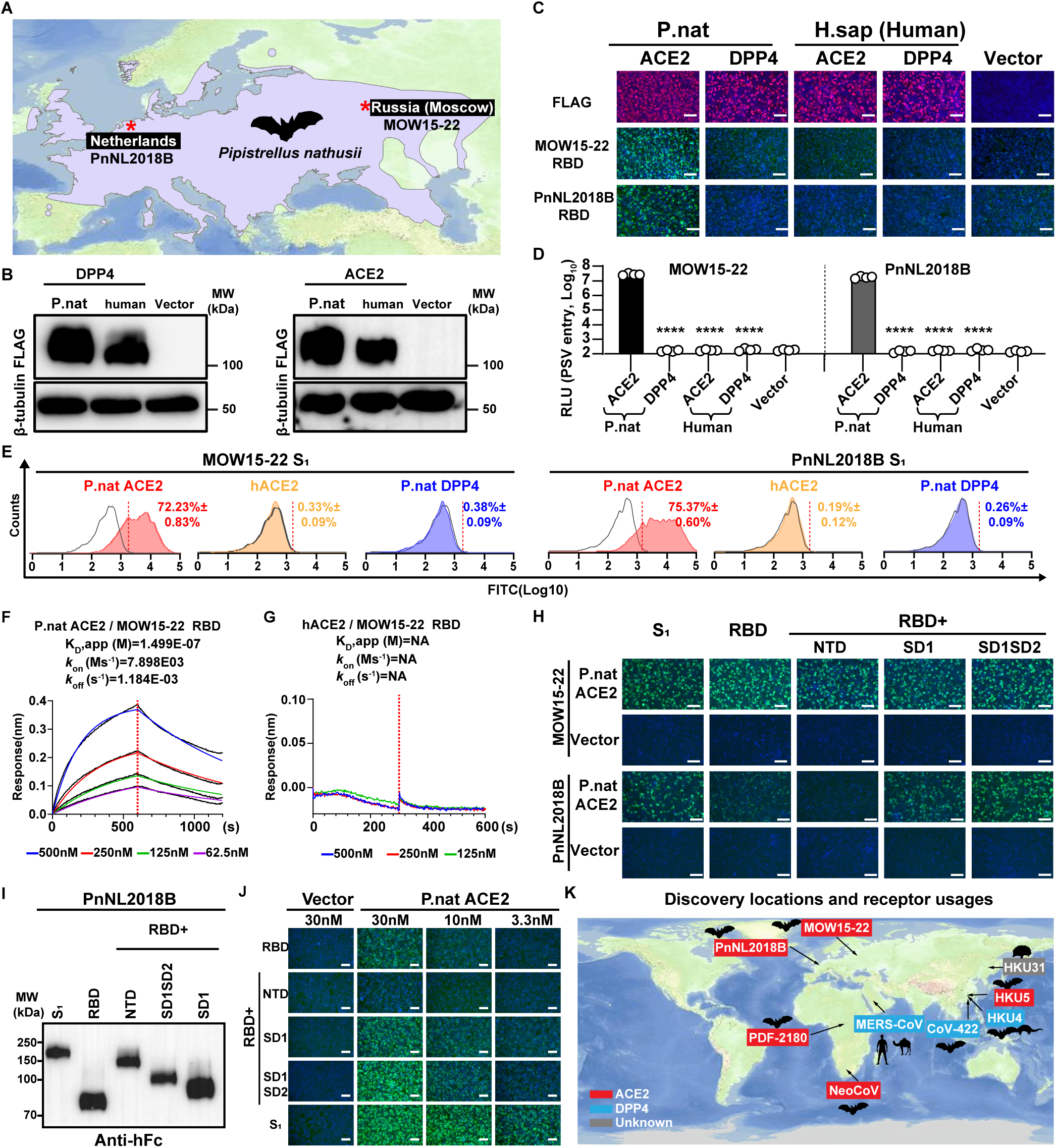
MOW15-22 and PnNL2018B use ACE2 as receptor. (**A**) Geographical distribution of *Pipistrellus nathusii* in Europe. The purple regions represent its habitat, data retrieved from the IUCN (International Union for Conservation of Nature) Red List of Threatened Species, and the distribution chart was generated using Geoscene Pro. Red asterisks indicate viral discovery locations. (B) Expression levels of human or P.nat ACE2 and DPP4 orthologs in HEK293T cells. (**C-D**) P.nat ACE2 but not DPP4 supports MOW15-22 and PnNL2018B RBD-hFc binding (C) and pseudovirus (PSV) entry (D) in HEK293T cells. Mean ± SD and unpaired two-tailed t-tests for D. n=4 biological replicates. RLU: Relative light unit. (**E**) Flow cytometry analysis of MOW15-22 and PnNL2018B S1 binding to P.nat ACE2, hACE2 or P.nat DPP4 transiently expressed in HEK293T cells. Gray: cells transfected with vector control. Mean ± SD for three technical repeats. (**F-G**) BLI analyses of binding kinetics of the soluble dimeric P.nat ACE2 (F) or hACE2 (G) ectodomain to immobilized MOW15-22 RBD-hFc. Analysis was conducted with curve-fitting kinetic with global fitting (1:1 binding model). NA: not applicable. (**H**) Binding of recombinant MOW15-22 and PnNL2018B S1 subunit constructs to P.nat ACE2 expressed at the surface of HEK293T cells detected by fluorescence microscopy. (**I**) Expression levels of recombinant proteins comprising different domains of the PnNL2018B S1 subunit. Equal volumes of protein-containing supernatants used for purification were loaded for Western blot analysis. (**J**) Binding of PnNL2018B S1 subunit constructs at various concentrations to HEK293T cells transiently expressing P.nat ACE2 analyzed by fluorescence microscopy. (**K**) Confirmed DPP4 or ACE2 usage of representative merbecoviruses is displayed with blue and red backgrounds, respectively. Discovery locations and natural hosts are indicated. Scale bars: 100 μm for C, H, and J.

However, in view of the unique sequence and structural features of MOW15-22 and PnNL2018B, we set out to investigate receptor usage of MOW15-22 and PnNL2018B by assessing the ability of human and P.nat ACE2 and DPP4 orthologs to support S-mediated pseudovirus entry and viral antigen bindings. We found that P.nat ACE2, but not human ACE2 (hACE2), human DPP4, or P.nat DPP4, promoted MOW15-22 and PnNL2018B pseudovirus entry and RBD or S1 binding (Figures 2B-2E). Biolayer interferometry (BLI) analysis showed that the soluble dimeric P.nat ectodomain, but not human ACE2, bound to immobilized MOW15-22 RBD with an apparent affinity (KD, app) of 149.9 nM (Figures 2F-2G). Further investigations of the contributions of the four domains present in the S1 subunit revealed that the MOW15-22 RBD is sufficient for binding to P.nat ACE2 whereas the SD1 and SD2 domains seem required for better PnNL2018B engagement (Figures 2H-2J). Therefore, we recommend using the S1 subunits of PnNL2018B to assess ACE2 binding efficiency.

Collectively, these data along with a recent report showing that HKU5 uses *Pipistrellus abramus* (P.abr) ACE2 as an entry receptor^32^ expand the geographic distribution of ACE2-using merbecoviruses to all three continents of the Old World, with Pipistrellus bats as important reservoir hosts that should be closely monitored (Figure 2K).

### Multi-species ACE2 tropism of MOW15-22 and PnNL2018B

To explore the potential host range of MOW15-22 and PnNL2018B, we assessed the ability of various ACE2 orthologs transiently transfected in HEK293T cells to promote RBD/S1 subunit binding and pseudovirus entry. We used a receptor library comprising 105 ACE2 orthologs from 52 bats and 53 non-bat mammalian species with validated expression (Figure S1).^24^ MOW15-22 efficiently used bat ACE2 from P.nat, P.par, P.dav, and a few others to a lesser extent (Figure 3A). PnNL2018B could only use P.nat and L.bor ACE2 efficiently for entry. In contrast to NeoCoV and PDF-2180, MOW15-22 and even more so PnNL2018B had limited tropism for non-bat mammalian ACE2s (Figure 3B), with only a few orthologs from Carnivora and Primates species, such as dog (C.fam) and common marmoset (C.jac) supporting MOW15-22 entry (Figure 3B).

**Figure 3.**
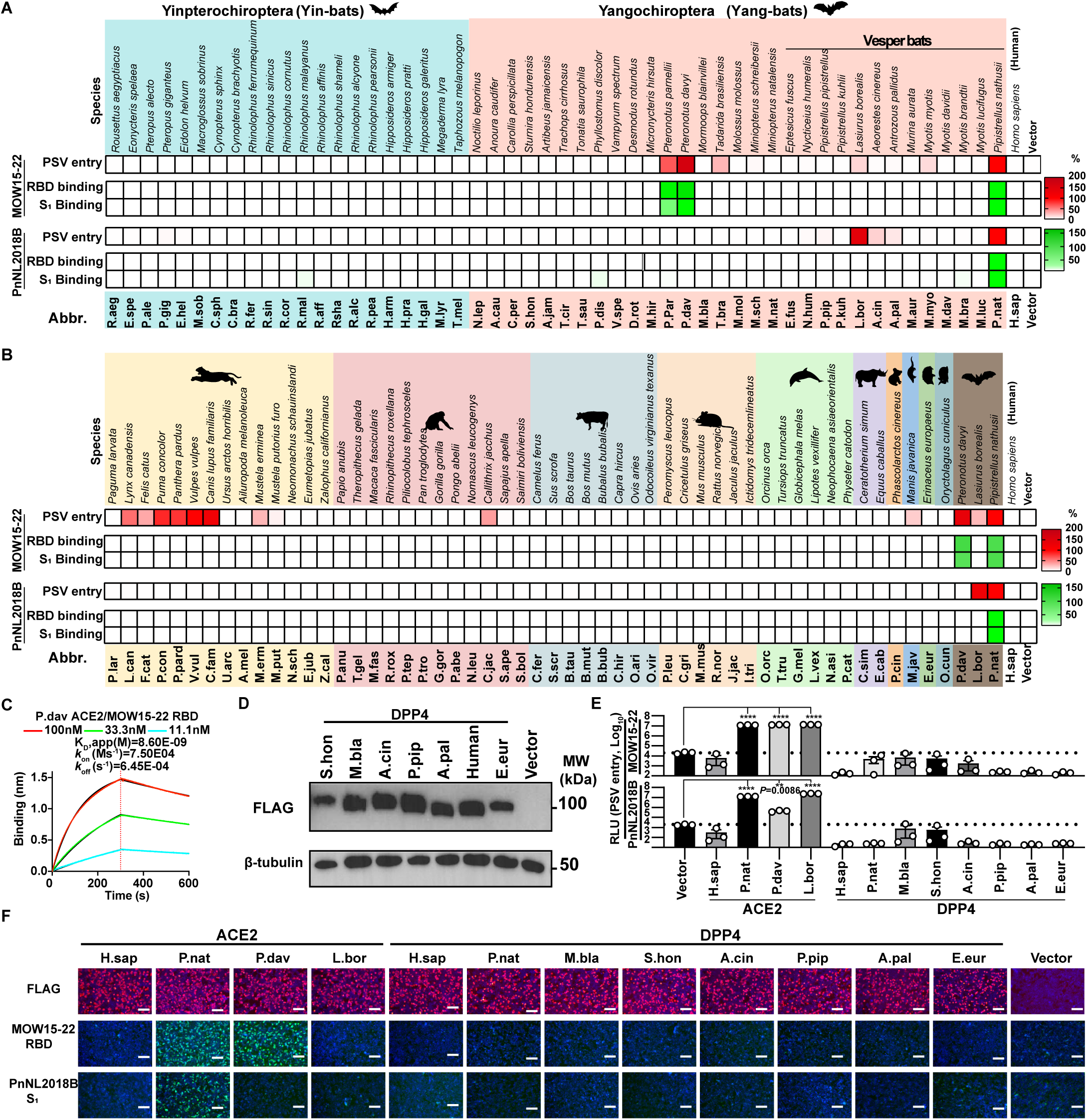
Multi-species ACE2 tropism of MOW15-22 and PnNL2018B. (**A-B**) Heat map representing MOW15-22 PSV entry into and RBD/S1 subunit binding to HEK293T cells transiently expressing various ACE2 orthologs from bats (A) or other mammalian species (B). Different mammalian orders are denoted with distinctly colored backgrounds, from left to right: Carnivora, Primates, Artiodactyla, Rodentia, Cetacea, Perissodactyla, Diprotodontia, Pholidota, Erinaceomorpha, and Lagomorpha. Data are normalized relative to P.nat ACE2 and shown as mean values. n=3 biological repeats for PSV entry. Data representative of 2-3 independent experiments for antigen binding assays. (**C**) BLI analysis of binding kinetics of the soluble recombinant dimeric P.dav ACE2 ectodomain to immobilized biotinylated MOW15-22 RBD. (**D**) Expression levels of several mammalian DPP4 orthologs in HEK293T cells. (**E**) PSV entry efficiency of MOW15-22 and PnNL2018B in HEK293T expressing the indicated receptors. Dashed lines: threshold of the background entry. MEAN ± SEM and unpaired two-tailed t-tests. n=3 biological replicates. (**F**) MOW15-22 and PnNL2018B RBD-hFc binding to HEK293T cells transiently expressing the indicated receptors assessed by fluorescence microscopy. Scale bars: 100 μm. See also Figure S1–S3.

Concurring with the entry data, P.nat, P.dav, P.par, and less prominent for several other mammalian ACE2s, supported binding of the MOW15-22 S1 subunit whereas only P.nat ACE2 supported efficient PnNL2018B RBD/S1 binding, with higher sensitivity observed in S1 binding assays (Figures 3A-3B and Figure S2). Conversely, ACE2 from Pipistrellus bats, P.pip and P.kuh, did not support MOW15-22 or PnNL2018B RBD/S1 binding, indicating host ACE2 specialization during virus evolution. As a result, two Pteronotus bat ACE2s but not the additional two Pipstrellus bat orthologs tested were functional receptors, underscoring the importance of specific sequence determinants over phylogenetic relationships of the bat hosts. BLI analysis showed that the soluble P.dav ACE2s bound to immobilized MOW15-22 RBD with an apparent affinity (KD, app) of 8.6 nM, outperforming the binding of P.nat ACE2 (Figure 3C). We also tested several DPP4 orthologs from bats, humans, and hedgehogs, and none of them showed any detectable receptor function (Figure 3D-F).

We observed pseudovirus entry for several ACE2 orthologs for which only weak or no MOW15-22 S1 binding could be detected (Figure 3B, Figure S2C, and S3A-B), as previously described for other coronaviruses.^9,33^ This apparent discrepancy may be attributed to the use of TPCK-treated trypsin to enhance pseudovirus entry with weak receptors and to the multivalent presentation of S trimers on pseudoviruses markedly increasing binding avidity, as compared to dimeric RBD or S1-Fc.^34^ Accordingly, L.bor ACE2 promoted pseudovirus entry and membrane fusion in the presence of exogenous trypsin, although no MOW15-22 S1 binding to this ACE2 was detected (Figure S3C-F). The addition of exogenous trypsin can thus improve pseudovirus entry assay sensitivity to identify weakly functional ACE2 orthologs.^34^

### Host ACE2 tropism determinants for MOW15-22 and PnNL2018B

We previously identified four host range molecular determinants (designated A-D) on the ACE2 receptor for NeoCoV/PDF-2180.^24^ To assess their relevance for species-specific recognition by MOW15-22 and PnNL2018B, we generated P.dav, P.par, P.pip, and L.bor ACE2 mutants harboring unfavorable changes for NeoCoV S-mediated entry. Unexpectedly, none of these alterations markedly affected MOW15-22 S receptor utilization (Figure S4). Moreover, exchanging residues 1-400 between M.bla ACE2 and P.dav ACE2 had no effect on their ability to promote MOW15-22 RBD binding and pseudovirus entry in spite of comprising all of the corresponding residues recognized by NeoCoV (Figure 4A). Additional sequence swaps led to the identification of residues 400-450 and subsequently the N428M.blaACE2 glycosylation site (Y430P.davACE2) as critical host range determinants for both MOW15-22 and PnNL2018B, with the presence of this oligosaccharide abolishing RBD or S1 binding with P.dav ACE2 (Figure 4A and Figure S5A-D). However, the restrictive effect of this glycan is less pronounced on MOW15-22/PnNL2018B S1 than MOW15-22/PnNL2018B RBD binding to P.nat ACE2, underscoring possible differences of interactions with different ACE2s (Figures S5E-5F). Furthermore, the N432C mutation, which abrogates the glycosylation site, was insufficient to enable strong binding of the MOW15-22 RBD or S1 to P.pip ACE2 without the additional substitutions of residues 500-600 or of residues E589P.pipACE2 (K in P.nat ACE2) and K597P.pipACE2 (E in P.nat ACE2) (Figure 4B). Consistently, P.nat ACE2 chimera with residues 500-600 replaced by P.pip ACE2 equivalents lost MOW15-22 RBD/S1 binding. Furthermore, substitutions at position K287P.natACE2 (K in P.nat ACE2) were identified to modulate MOW15- 22 and PnNL2018B RBD and S1 binding in ACE2 swap experiments testing P.nat ACE2 and P.pip ACE2-N432C (Figure 4C). Given that the residues identified as governing receptor utilization are spatially close to each other but far from the NeoCoV/PDF-2180 footprint on ACE2 along with their different phenotypes of sensitivity to specific ACE2 glycans, we hypothesized that MOW15-22 and PnNL2018B might rely on a distinct mode of receptor engagement (Figure 4D-E).

**Figure 4.**
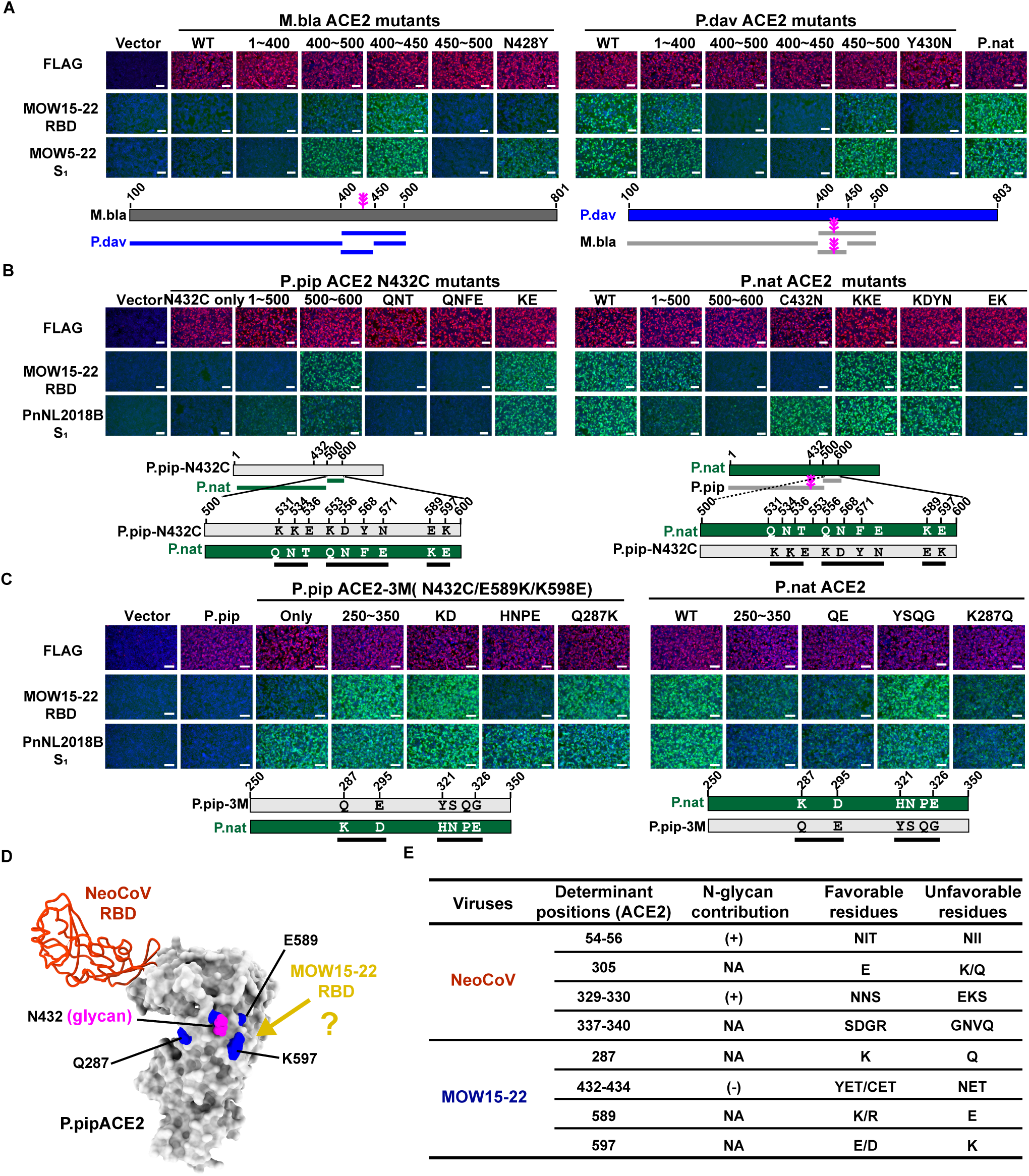
Host ACE2 tropism determinants for MOW15-22 and PnNL2018B. (**A**) MOW15- 22 RBD and S1 subunit binding to HEK293T cells transiently expressing ACE2 chimeras with sequence swaps between M.bla and P.dav ACE2s analyzed by fluorescence microscopy. (**B**) Sequence swaps between P.pip ACE2 N432C mutant and P.nat ACE2 and their impact on indicated viral antigen bindings. (**C**) Sequence swaps between P.pip ACE2 N432C/E589K/K598E mutant and P.nat ACE2 and their impact on indicated viral antigen bindings. (**D**) Summary of the critical determinants for MOW15-22 receptor recognition and host range determination identified through genetic swaps. Key residues discussed in the text are highlighted in blue on the P.pip ACE2 surface (PDB 7WPO). The N432-glycan is indicated in magenta. (**E**) Summary of ACE2 residues and glycans governing receptor utilization for MOW15-22 or PnNL2018B, respectively. The contribution of N-glycans in receptor binding and representative favorable/unfavorable residues in ACE2 orthologs are indicated. (**+**): support binding; (**-**): restrict binding; NA: not applicable (not glycosylation site). The P.pip ACE2 residue numbering is shown. Residue positions, N432P.pip ACE2 glycans (in magenta), and swap strategies were indicated with the schematic below each panel in A, B, and C. Scale bars:100 μm. See also Figure S4–S5.

### Structural basis for MOW15-22 binding with bat ACE2

To understand the molecular basis of ACE2 utilization by these recently identified merbecoviruses, we determined a cryo-EM structure of the MOW15-22 RBD bound to the dimeric P.dav ACE2 ectodomain at 2.8Å resolution (Figure 5A, Figure S6, and Table S1). Two MOW15-22 RBD extensions spanning residues 505-535 and 563-571 protrude at the distal end of the receptor-binding motif (RBM) to interact with the ACE2 peptidase domain (Figure 5A). Specifically, the MOW15-22 RBM interacts with a glycan-free P.dav ACE2 surface comprising residues P282, Y283, E285, K286, P287, P427, E428, D429, Y430, E431, E433, I434, L437, T534, P536, H538, R587, P588, N591, W592, E595, Q596 and S598, encompassing the identified host range-associated P.dav ACE2 residues (E285, Y430, R587, and E595) (Figure 4D). An average surface of ∼850Å^2^ is buried at the interface between the RBD and ACE2 with key interactions including (Figure 5B-C): (**i**) K509MOW15-22 forming a salt bridge with E595ACE2; (**ii**) a salt bridge triad involving K286ACE2 with D563MOW15-22 and D567MOW15-22; (**iii**) a hydrogen bond network between Y430ACE2, T526MOW15-22, and R571MOW15-22; (**iv**) hydrogen bonding between N591ACE2 and E523MOW15-22; (**v**) Y283ACE2 that is hydrogen-bonded to the D508MOW15-22 backbone carbonyl and tucked in between V507MOW15-22 and P511MOW15-22; and (**vi**) extensive hydrophobic interactions mediated by V507MOW15-22, I521MOW15-22, and F524MOW15-22 with ACE2.

**Figure 5.**
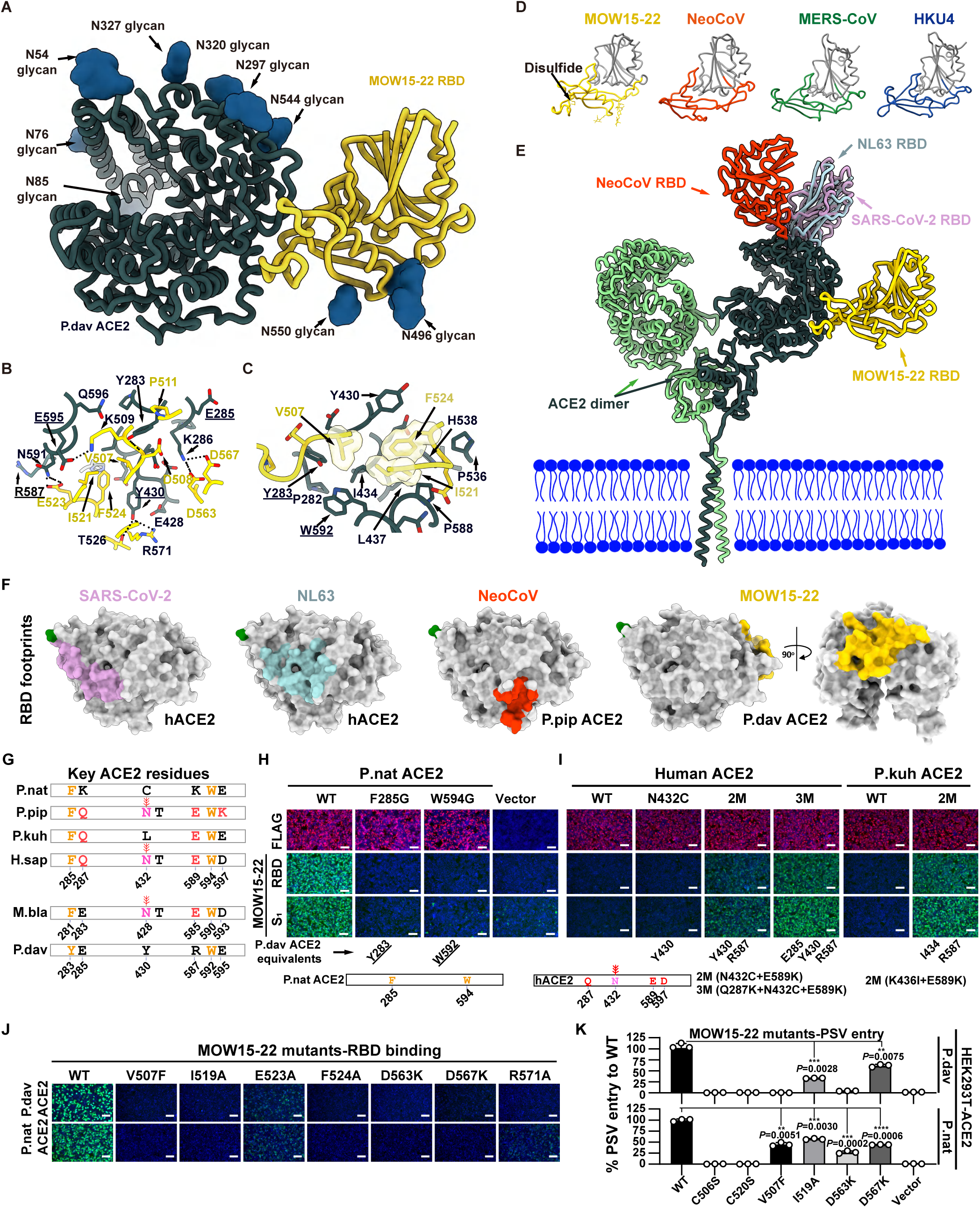
Structural basis for MOW15-22 binding with bat ACE2. (**A**) CryoEM structure of the MOW15-22 RBD bound to the P. dav ACE2 ectodomain. (**B-C**) Zoomed-in views of key interactions mediating MOW15-22 RBD(yellow) binding to P.dav ACE2 (dark green). Selected salt bridges and hydrogen bonds are shown as black dotted lines. (**D**) Comparison of merbecovirus RBD architectures. MOW15-22, MERS-CoV (PDB 6Q04), HKU4 (PDB 4QZV), NeoCoV (PDB 7WPO). RBD core subdomains are indicated in gray. (**E)** Comparison of the binding modes of the MOW15-22, NeoCoV (PDB 7WPO), SARS-CoV-2/SARS-CoV-1 (PDB 7TN0), and NL63 (PDB 3KBH) RBDs on bat (not shown for clarity) or hACE2 (PDB 6M1D, B0AT1 not shown for clarity). (**F**) RBD footprints of the four ACE2-using coronaviruses. N- terminus labeled in green. (**G**) Schematic of critical residues responsible for species-specific receptor recognition in selected ACE2 orthologs. (**H**) P. nat ACE2 mutants with reduced MOW15-22 RBD or S1 subunit binding due to unfavorable substitutions to hydrophobic interactions. (**I**) human or P.kuh ACE2 mutants with improved MOW15-22 RBD or S1 subunit binding due to favorable substitutions mapping to the N432-glycosylation site, hydrophobic interactions, or polar interactions. (**J-K**) MOW15-22 RBM mutants with reduced ability to utilize P.dav and P. nat ACE2 orthologs for RBD binding (J) and pseudovirus entry (K). Scale bars:100 μm. Binding in panels H, I, and J was analyzed by fluorescence microscopy. MEAN ± SD and unpaired two-tailed t-tests for K. n=3 biological replicates. See also Figure S6–S7.

The MOW15-22 RBD largely adopts a canonical merbecovirus architecture comprising a core folded as a five-stranded antiparallel β-sheet with two α-helices and a four-stranded antiparallel β-sheet RBM with one α-helix (Figure 5D). However, the MOW15-22 RBD is set apart from other merbecoviruses due to its elongated and twisted distal RBM extensions, one of them being stabilized by the C506-C520 disulfide, which form the ACE2-binding surface, and to the elongated α-helix flanking the RBM on the opposite side. Furthermore, the MOW15-22 RBD harbors two N-linked glycans protruding from the base of the RBM, which are conserved in PnNL2018B S but not among other merbecoviruses (Figure 5D). The reduced P.dav ACE2 binding and utilization observed for the PnNL2018B RBD, relative to MOW15-22, is explained by the conservation of 14 out of 21 ACE2-interacting residues (Figure 5D and Table S2).

Strikingly, the MOW15-22 RBD recognizes ACE2 at a site located more than 50Å away from the sites recognized by SARS-CoV-2, NL63, or NeoCoV (Figures 5E-5F). This suggests that ACE2-utilization was acquired multiple times independently among these coronaviruses, including at least twice for MERSr-CoVs. Our data reveal that P.dav ACE2 lacks a glycan at position Y430, which is otherwise conserved in several other ACE2 orthologs (e.g., hACE2/P.pip N432 and M.bla N428 glycans), and would impede MOW15-22 RBD binding sterically, concurring with the lack of binding observed to a P.dav Y430N glycan knockin mutant (Figure 5C). Accordingly, ACE2 orthologs lacked this glycan (e.g. P. par ACE2, C.jac, and L.bor ACE2) supporting efficient MOW15-22 or PnNL2018B S pseudovirus entry into cells (Figures 3A-3B), and these viruses are therefore not expected to acquire efficient hACE2 utilization without major adaptations.

To further assess the contribution of the identified receptor binding determinants, we analyzed the impact of residue changes at key positions of P.nat, P.pip, human, P.kuh, and C.jac orthologs (Figure 5G). P.nat ACE2 mutants harboring F285G and W594G mutations with reduced hydrophobicity dampened binding to MOW15-22 or PnNL2018B, while three hACE2 residue substitutions (Q287K, N432C, and E589K) and two P.kuh ACE2 residue substitutions (K436I and E589K) promoted detectable binding of MOW15-22 and PnNL2018B (Figures 5H 5I and Figure S7A-7B). The C.jac ACE2 S432N glycan knock-in mutation abolished receptor function of the sole primate ortholog supporting entry, further underscoring the key role of this oligosaccharide as a tropism barrier. Conversely, the C.jac ACE2 Q287K and Q598L point mutations with additional interactions significantly improved binding affinity to both MOW15-22 and PnNL2018B (Figure S7C-7D). Concurring with the structural data, introducing unfavorable changes to the MOW15-22 RBM, either by reducing interaction or abolishing the C506-C520 disulfide, significantly reduced RBD binding and pseudovirus entry in P.nat or P.dav ACE2-expressing HEK293T cells (Figures 5J-5K and Figure S7E).

### Characterization and inhibition of MOW15-22 and PnNL2018B ACE2-mediated entry

MERS-CoV efficiently utilizes endogenous host proteases, such as furin and TMPRSS2, for cellular entry.^35^ MERS-CoV S harbors a polybasic (furin) cleavage site at the junction between the S1/S2 subunits, leading to proteolytic processing during biogenesis.^36^ Conversely, MOW15- 22 and PnNL2018B S glycoproteins lack an S1/S2 furin cleavage site and remain uncleaved upon incorporation into VSV pseudovirus particles, suggesting distinct protease requirements for cellular entry compared to MERS-CoV (Figures 6A-B). Accordingly, MOW15-22 and PnNL2018B S glycoproteins are highly dependent on the addition of exogenous trypsin to promote cell-cell fusion, indicating a potential reliance on endocytic pathways for cellular entry in Caco2 cells, which endogenously express TMPRSS2 at their surface (Figures 6C-6D).^37^

**Figure 6.**
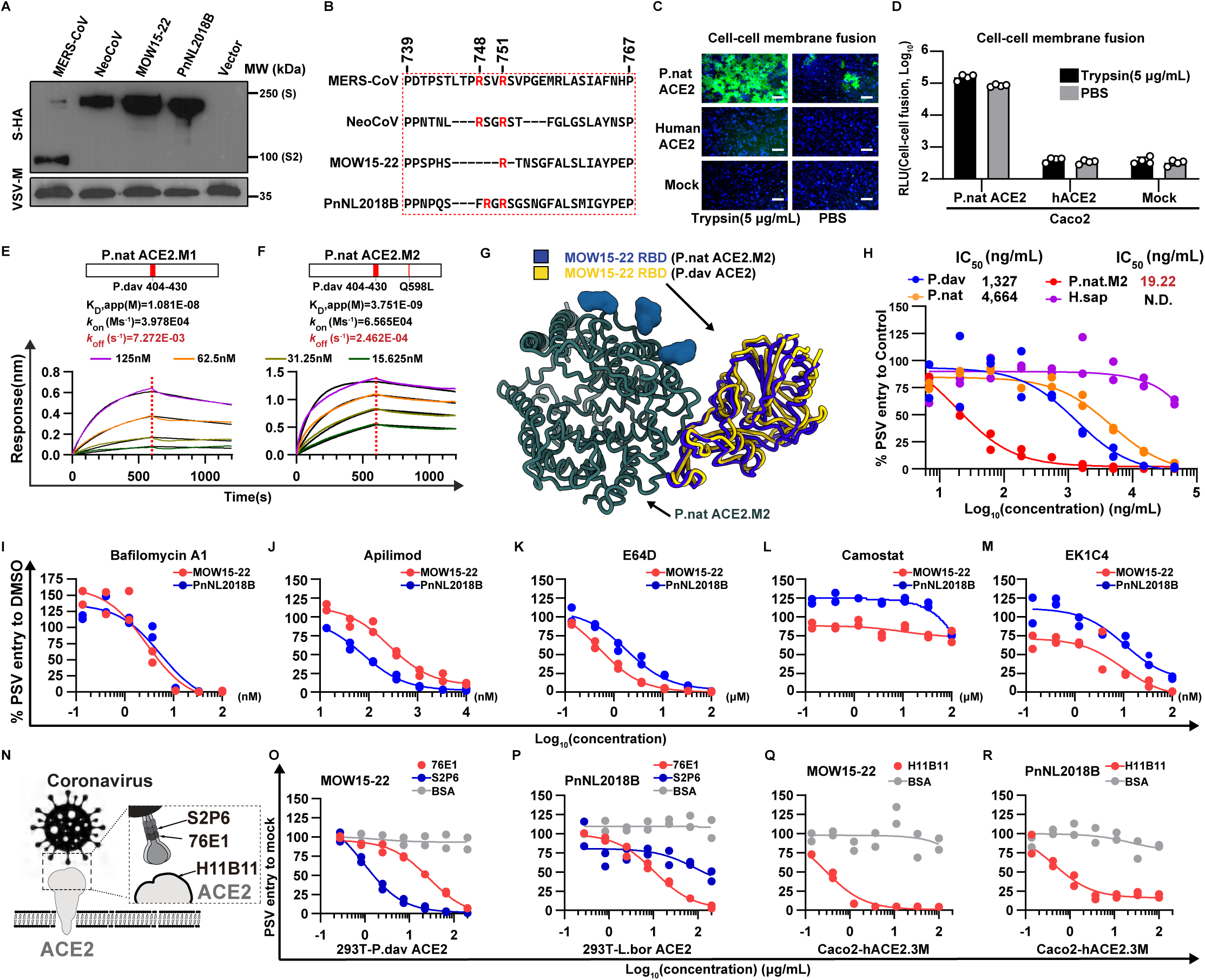
Characterization and inhibition of MOW15-22 and PnNL2018B ACE2-mediated entry. (**A**) Quantification of S glycoprotein incorporation in VSV pseudotypes analyzed by Western blot detecting the C-terminal-fused HA tags. VSV-M was used as a loading control. (**B**) Sequence analysis of S glycoprotein S1/S2 junction (dashed box) from the indicated viruses, with arginine highlighted in red. (**C-D**) MOW15-22 S mediated cell-cell fusion using Caco2 cells overexpressing P.nat ACE2 upon addition of TPCK-treated trypsin. The signal resulting from the reconstitution of split sfGFP (C) or RLuc (D) is indicated. (**E-F**) BLI analyses of binding kinetics of soluble dimeric ACE2 ectodomains from P.nat ACE2.M1 (E) and P.nat ACE2.M2 (F) carrying P.dav ACE2 equivalent residues (red) to immobilized MOW15-22 RBD-hFc. Global fitting to the data using a 1:1 binding model is shown in black. (**G**) Superimposition of the cryoEM structures of the MOW15-22 RBD (gold) in complex with P.dav ACE2 (not shown for clarity) and of the MOW15-22 RBD (blue) bound to P.nat ACE2.M2 (dark green) superimposed based on the ACE2 peptidase domain. (**H**) Dose-dependent inhibition of MOW15-22 pseudovirus entry by soluble P.dav, P.nat, P.nat.M2 and human ACE2s in Caco2 cells stably expressing P.nat ACE2. IC50 values are indicated. N.D.: Not detectable. (**I-M**) Inhibitory activities of small compounds or peptide inhibitors against MOW15-22 and PnNL2018B pseudoviruses using HEK293T cells stably expressing P.dav ACE2 (for MOW15-22) or L.bor ACE2 (for PnNL2018B). Endosomal acidification inhibitor bafilomycin A1 (Baf-A1) (I), and the PIKfyve inhibitor Apilimod (J), cathepsin L inhibitor E64d (K), TMPRSS2 inhibitor camostat (L), S2-targeting HR1 peptide fusion inhibitor EK1C4 (M). (**N**) Schematic representation of the epitopes targeted by the antibodies used in the pseudovirus neutralization assays. (**O-R**) Dose- dependent inhibition of MOW15-22 and PnNL2018B pseudoviruses by S2P6 and 76E1 (O-P), or H11B11 (Q-R) in the indicated cell lines.n=2 biological replicates for H-R. Scale bars:200 μm. See also Figure S8–S10.

To compete viral entry using soluble ACE2, we designed P.nat ACE2 recombinant ectodomain chimeras carrying P. dav ACE2 residues 404-432 without (P. nat ACE2.M1) or with the Q598L substitution (P.nat ACE2.M2) which exhibited enhanced binding affinity for both MOW15-22 and PnNL2018B RBD-Fc constructs relative to wildtype P. nat ACE2, especially the off-rate that is most critical for neutralization (Figures 6E-6F, Figure S8A-C).^9^ Cryo-EM analysis of the complex between the MOW15-22 RBD and P. nat ACE2.M2 confirmed the retention of a native binding mode and rationalized the enhanced binding (Figure 6G, Figure S9, and Table S1). We observed concentration-dependent soluble P.nat ACE2.M2-mediated inhibition of MOW15-22 and PnNL2018B S pseudoviruses with a half-maximal inhibitory concentration (IC50) of 19.22 ng/ml, outperforming wild-type P.nat and P.dav ACE2s (Figure 6H), further confirming the potential of these ACE2 mutants for developing entry inhibitors.

To identify other countermeasures against these divergent merbecoviruses, we assessed the ability of entry inhibitors, monoclonal antibodies, nanobodies, and peptide fusion inhibitors to block S-mediated entry in HEK293T cells expressing P.dav ACE2 (Figures 6I-6R and Figure S10). Endosomal entry inhibitors, such as E64D, bafilomycin A, and Apilimod, inhibited pseudovirus entry in a dose-dependent manner (Figures 6I-6K) whereas the TMPRSS2 inhibitor camostat did not, likely due to the lack of TMPRSS2 expression in this cell line or their low efficiency in using this protease (Figure 6L).^37–39^ These results confirmed that MOW15-22 S- and PnNL2018B S-mediated entry can occur through an endosomal pathway in these target cell types. Whereas MERS-CoV RBD-directed nanobodies were ineffective against MOW15-22 and PnNL2018B S pseudoviruses (Figure S10A),^40,41^ broadly neutralizing S2 subunit-directed monoclonal antibodies targeting the stem helix (S2P6) or the fusion peptide/S2’ cleavage site (76E1) retained activity, consistent with the conservation of the targeted S2 epitopes and the marked divergence of their RBDs (Figures 6N-6R and Figures S10B-D).^42,43^ Furthermore, the HR1-directed EK1C4 peptide and the hACE2-targeting antibody H11B11 efficiently neutralized the entry of MOW15-22 and PnNL2018B pseudoviruses (Figures 6M, Q, R and Figure S10E).^39,44^ Collectively, these data demonstrate that neutralizing antibodies targeting the fusion machinery (S2 subunit) or the host receptor prevent ACE2-mediated entry and could be potentially used as countermeasures against these viruses.

### MOW15-22 and PnNL2018B S-mediated propagation supported by ACE2

Finally, we utilized a reverse genetic system to create propagation-competent VSV (pcVSV) to characterize the MOW15-22 and PnNL2018B S glycoprotein-mediated propagation supported by different ACE2. The VSV-G genes in these recombinant viruses were replaced with the MOW15-22 or the PnNL2018B S glycoprotein gene, with an additional GFP gene for visualization (Figure 7A-7B). We successfully rescued the viruses and observed a dose- dependent trypsin enhancement of amplification in Caco2 cells expressing P.nat ACE2, as evidenced by syncytia formation (Figure 7C). Although viral propagation was not observed in Caco2 cells overexpressing hACE2, abrogation of the N432 glycan (via the N432C mutation) resulted in detectable amplification of pcVSV-MOW15-22. Furthermore, the E589K or Q287K/E589K mutations in the hACE2 N432C background further improved the amplification of pcVSV-MOW15-22 (Figure 7D) and enabled the detection of pcVSV-PnNL2018B amplification (Figure 7E), concurring with our binding data (Figure 5H). Despite the restriction on viral recognition imposed by the hACE2 N432 glycan, the two viruses could nevertheless utilize P.nat ACE2 harboring a C432N (glycan knockin) mutation, as demonstrated by the observed viral propagation and S1 subunit binding (Figures 7F-7G and Figure S5E-F). The observed phenotype indicates that these viruses hold the potential to partially overcome the steric tropism barrier imposed by the N432 glycan, possibly through compensatory interactions in other ACE2 regions.

**Figure 7.**
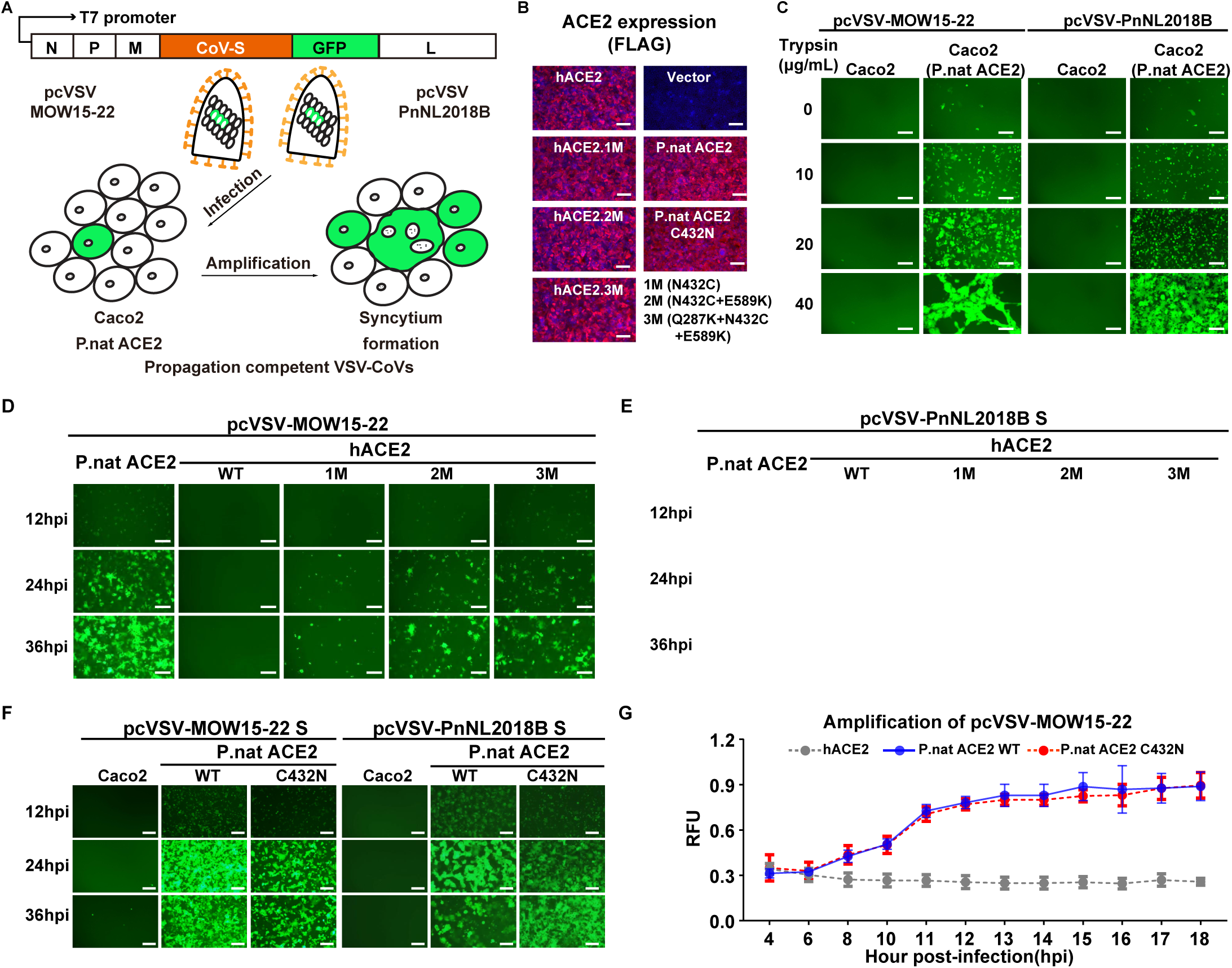
MOW15-22 and PnNL2018B S-mediated propagation supported by ACE2. (**A**) Genetic organization and amplification of pcVSV-MOW15-22 and pcVSV-PnNL2018B. (**B**) Immunofluorescence analyzing the expression of P.nat and human ACE2 mutants stably expressing in Caco2 cells. (**C**) GFP signal from TPCK-trypsin enhanced amplification of pcVSV-MOW15-22 and pcVSV-PnNL2018B in Caco2 cells with or without the expression of P.nat ACE2 at 24hpi. (**D-E**) Propagation of pcVSV-MOW15-22(D) and pcVSV-PnNL2018B(E) in Caco2 cells stably expressing hACE2 carrying N432C(1M), Q287K/N432C(2M), Q287K/N432C/E589K(3M) mutants, with P.nat ACE2 as a positive control. (**F**) Representative fluorescence images of pcVSV-MOW15-22 and pcVSV-PnNL2018B amplification in Caco2 cells expressing P.nat ACE2 or P.nat ACE2 C432N mutants at 12, 24 and 36-hour post-infection (hpi). (**G**) pcVSV-MOW15-22 propagation kinetics in Caco2 cells expressing P.nat ACE2 or P.nat ACE2 C432N mutants at the indicated time points. MEAN ± SD and n=3 biological replicates. RFU: relative fluorescence unit. TPCK-treated trypsin was present in the cells at 20 μg/mL in D, E, F, and G. Scale bars:100 μm for B and 200 μm for C-E.

## DISCUSSION

Coronaviruses exhibit remarkable variations in RBD sequences, resulting in diverse receptor usage across different viruses.^45^ While coronaviruses within the same genus or subgenus typically share similar RBD core structures, RBM variability can result in entirely different receptor usage.^9,46,47^ Conversely, phylogenetically distant coronaviruses can convergently evolve to engage the same receptor during evolution. For example, APN is a receptor shared by many α- coronaviruses and δ-coronaviruses,^47–50^ whereas ACE2 serves as a functional receptor for three viral subgenera that belong to α-coronaviruses (*Setracoviruses*) and β-coronaviruses (*Sarbecoviruses* and *Merbecoviruses*) .^21,51–53^

The discovery of ACE2 usage in European bat MERSr-CoVs expands the diversity of ACE2-using merbecoviruses beyond NeoCoV and PDF-2180 and reveals that coronaviruses belonging to the same subgenus or even species (e.g. MOW15-22 and NeoCoV) can engage entirely distinct surfaces of a same receptor (e.g. P.dav ACE2). This study emphasizes the need to characterize receptor usage and binding modes experimentally, instead of solely relying on *in silico* predictions.

Our data further support the hypothesis that ACE2 usage was convergently opted by different coronaviruses with remarkable differences in their RBD or RBM sequences and structures. Furthermore, a recent report showed that HKU5 exhibited a specialized adaptation to its host ACE2 (P.abr), suggesting a potential fifth ACE2 binding mode that remains to be elucidated.^32^ Notably, these ACE2-using merbecoviruses with distinct binding modes evolved in three different Old World continents, suggesting the convergent evolution of ACE2 adaptation and the importance of investigating the global prevalence and distribution of ACE2-using merbecoviruses. This ACE2 preference likely confers certain evolutionary advantages in transmission, as exemplified by the highly transmissible SARS-CoV-2 omicron variants.^54,55^ However, a recent study reported that PnNL2180B (PN-βCoV) primarily exhibits intestinal tropism in its natural bat host, suggesting a potential fecal-oral route used by these viruses in bats .^28^ Given that airborne transmission is the major route of all known ACE2-using human coronaviruses, it is important to investigate whether tissue tropism and transmission route changes when ACE2-using viruses jump from bats to other mammals, especially humans.

Interestingly, ACE2 glycans can play contrasting roles in host range determination of ACE2-using merbecoviruses. For example, unlike the positive role of ACE2 glycans (e.g. N54 and N329 glycans) in NeoCoV and PDF-2180 recognition^9^, the N432 glycan in interaction interface restricted efficient MOW15-22 and PnNL2018B binding, contributing to narrower ACE2 tropism, particularly for PnNL2018B for which we only detected binding to its host P. nat ACE2^9^. Common marmoset (C.jac) ACE2, which is the only primate ortholog supporting MOW15-22 S-mediated entry, lacks an N432 glycan which is otherwise conserved in other primate ACE2s, explaining its distinct phenotypes. This specialized adaptation to host ACE2s lacking an N432-glycan is expected to prevent ACE2 utilization from many other species, including humans. However, the efficient amplification of pcVSV-MOW15-22 and pcVSV- PnNL2018B supported by a P.nat ACE2 mutant (C432N) harboring an N432 glycan suggests putative evolutionary pathways by which these viruses could overcome this obstacle and the usefulness of the countermeasures identified here. Notably, the VM314 viral sequence, identified in samples collected in the Netherlands in 2008 based on an RNA-dependent RNA polymerase (RdRp) gene fragment that is phylogenetically close to MOW15-22 and PnNL2018B, may already be adapted to its host (P.pip) ACE2 carrying an N432 glycan. Future studies should be conducted to obtain the corresponding S glycoprotein sequences and to verify whether these viruses recognize ACE2 in a similar way to MOW15-22.^56^

Previous studies proposed that the emergence of the DPP4-using MERS-CoV may be associated with a receptor switch from ACE2 to DPP4 through recombination.^9^ This could have occurred between an ACE2-using bat MERSr-CoV and a yet-to-be-identified DPP4-using merbecovirus, such as HKU4 or CoV-422.^15,57^ We note that CoV-422 was also suggested to have originated from recombinations leading to receptor switch^15^. Additionally, it has been suggested that ACE2-using merbecoviruses might have arisen through recombination between ancestral viruses of bats and hedgehogs.^58^ However, thus far, we and others have not detected solid evidence of ACE2 usage in testing bat or hedgehog ACE2 for coronaviruses HKU31 or other EriCoVs.^9,59,60^ These observations raise intriguing and important questions regarding the evolution trajectory of receptor usages of merbecoviruses and whether ACE2 receptor usage is the more ancestral trait for merbecoviruses than DPP4.

Our study profoundly changes our understanding of ACE2-using merbecoviruses by identifying and characterizing two bat merbecoviruses with previously unknown ACE2 binding modes, involving binding sites located 50Å away from that recognized by any other coronaviruses. This discovery also underscores the likelihood of the existence of other yet-to-be- discovered ACE2-using merbecoviruses, further expanding the diversity and geographic distribution of these viruses with spillover potential. Although the pathogenicity and transmission abilities of these viruses remain unclear, enhanced surveillance along with identification of viral inhibitors are warranted to proactively detect and prepare for potential zoonosis caused by ACE2-using merbecoviruses.

## Acknowledgments

This study was supported by the National Key R&D Program of China (2023YFC2605500 to H.Y. and Z.L.S.), the National Natural Science Foundation of China (NSFC) projects (82322041, 32270164, 32070160, 32188101 to H.Y., 323B2006 to C.B.M.), the Fundamental Research Funds for the Central Universities (2042023kf0191, 2042022kf1188 to H.Y.), Natural Science Foundation of Hubei Province (2023AFA015 to H.Y.), the National Institute of Allergy and Infectious Diseases (P01AI167966, DP1AI158186 and 75N93022C00036 to D.V.), an Investigators in the Pathogenesis of Infectious Disease Awards from the Burroughs Wellcome Fund (D.V.), the University of Washington Arnold and Mabel Beckman cryoEM center and the National Institute of Health grant S10OD032290 (to D.V.). D.V. is an Investigator of the Howard Hughes Medical Institute and the Hans Neurath Endowed Chair in Biochemistry at the University of Washington. The CAS Pioneer Hundred Talents Program to Z.D.

We are grateful to Michael Hiller (LOEWE Centre for Translational Biodiversity Genomics), Camila Mazzoni (Leibniz Institute for Zoo and Wildlife), and Huabin Zhao (Wuhan University) for sharing the ACE2 and DPP4 coding sequences of *Pipistrellus nathusii*. We thank Qiang Ding (Tsinghua University) and Qihui Wang (CAS Key Laboratory of Pathogenic Microbiology & Immunology, China) for sharing some mammalian ACE2 plasmids. We thank Mingzhou Chen (Hubei University) for providing the recombinant vaccinia virus expressing T7 RNA polymerase (vvT7). We thank the Center for Instrumental Analysis and Metrology of Wuhan Institute of Virology for providing technical assistance in cryoEM sample preparation and data collection, as well as the core facilities of the Key Laboratory of Virology, Wuhan University for providing instrumental and technical support.

## Author contributions

H.Y., Y.J.P., M.A.T., D.V., Z.D., Z.L.S, C.B.M., C.L., and Q.X., conceived the project; J.C., C.B.M., and C.L. conducted bioinformatic analysis; C.B.M. and C.L., designed and carried out experiments to generate clones, purify proteins, dissect host range determinants and test entry inhibitors. C.B.M., C.L., Y.J.P, Q.X., X.Yu., P.L., J.Y.S., M.L.H, F.T., Y.H.S., and Y.M.C. established assays and methods used in this study; X.Yang rescued the pcVSV-MOW15-22 and pcVSV-PnNL2018B viruses. J.L., J.B., and C.S. produced recombinant glycoproteins and carried out binding and entry assays. Y.J.P. carried out cryo-EM sample preparation, data collection, and processing of the P.dav ACE2-MOW15-22 RBD complex. D.A. helped with data collection. Y.J.P. and D.V. built and refined the P.dav ACE2-bound and MOW15-22 RBD structure; J.T. carried out cryo-EM sample preparation, data collection, and processing of the P.nat ACE2.M2-MOW15-22 RBD complex; Z.D. and J.T. built and refined the P.nat.ACE2.M2 ACE2-MOW15-22 RBD complex structure. H.Y., and D.V., wrote an initial draft of the manuscript with input from all authors.

## STAR ★ METHODS

### KEY RESOURCES TABLE

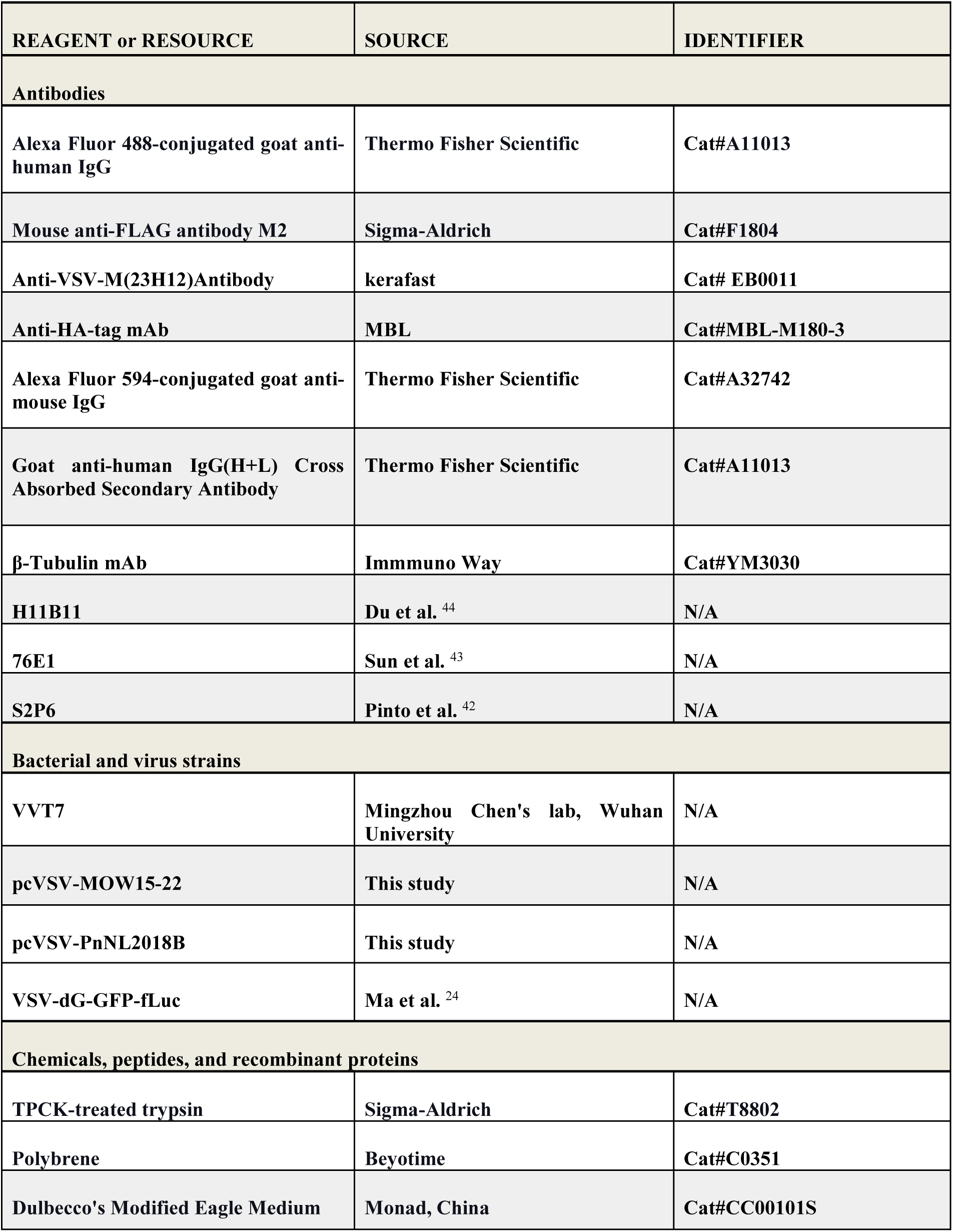
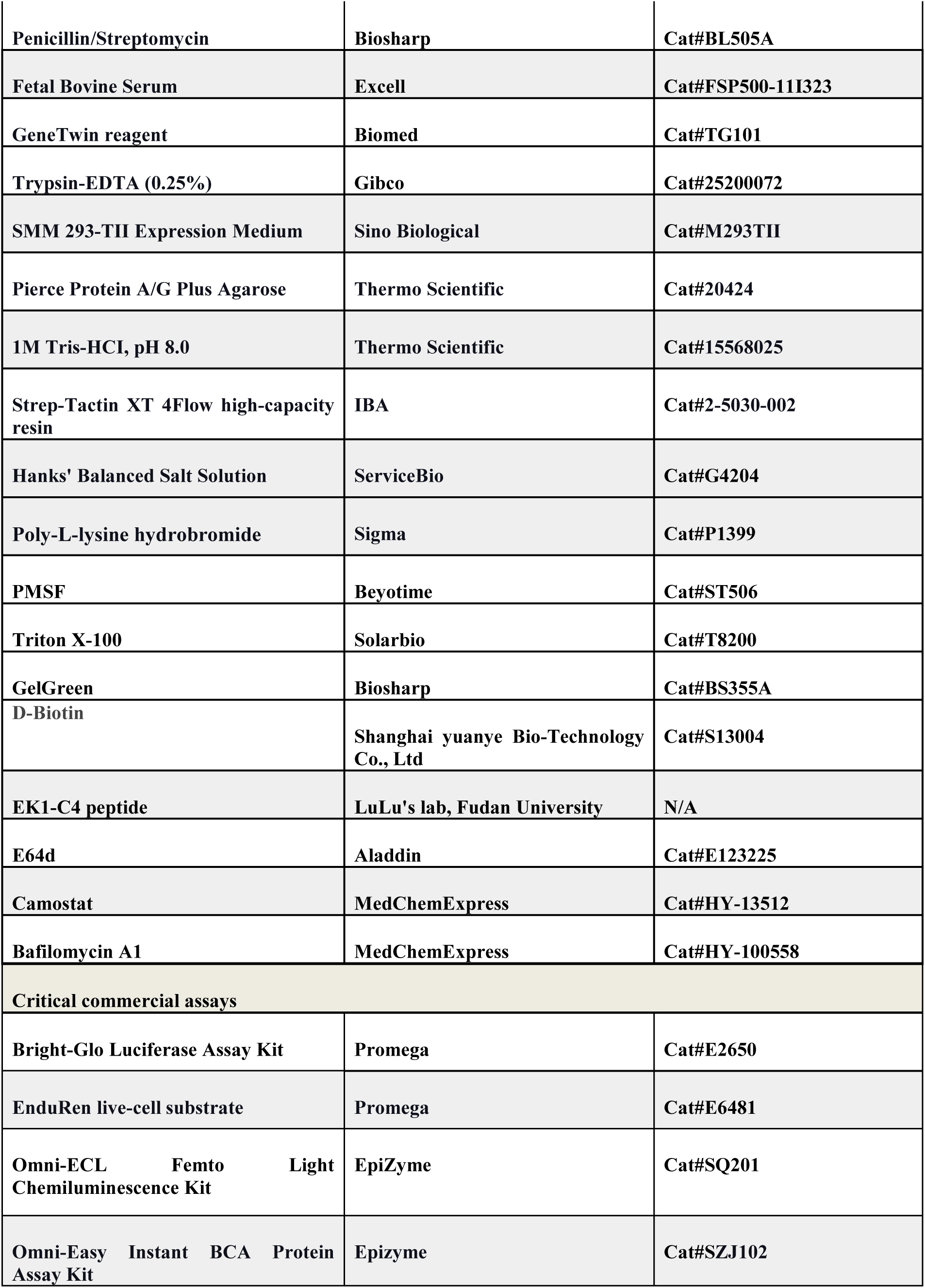
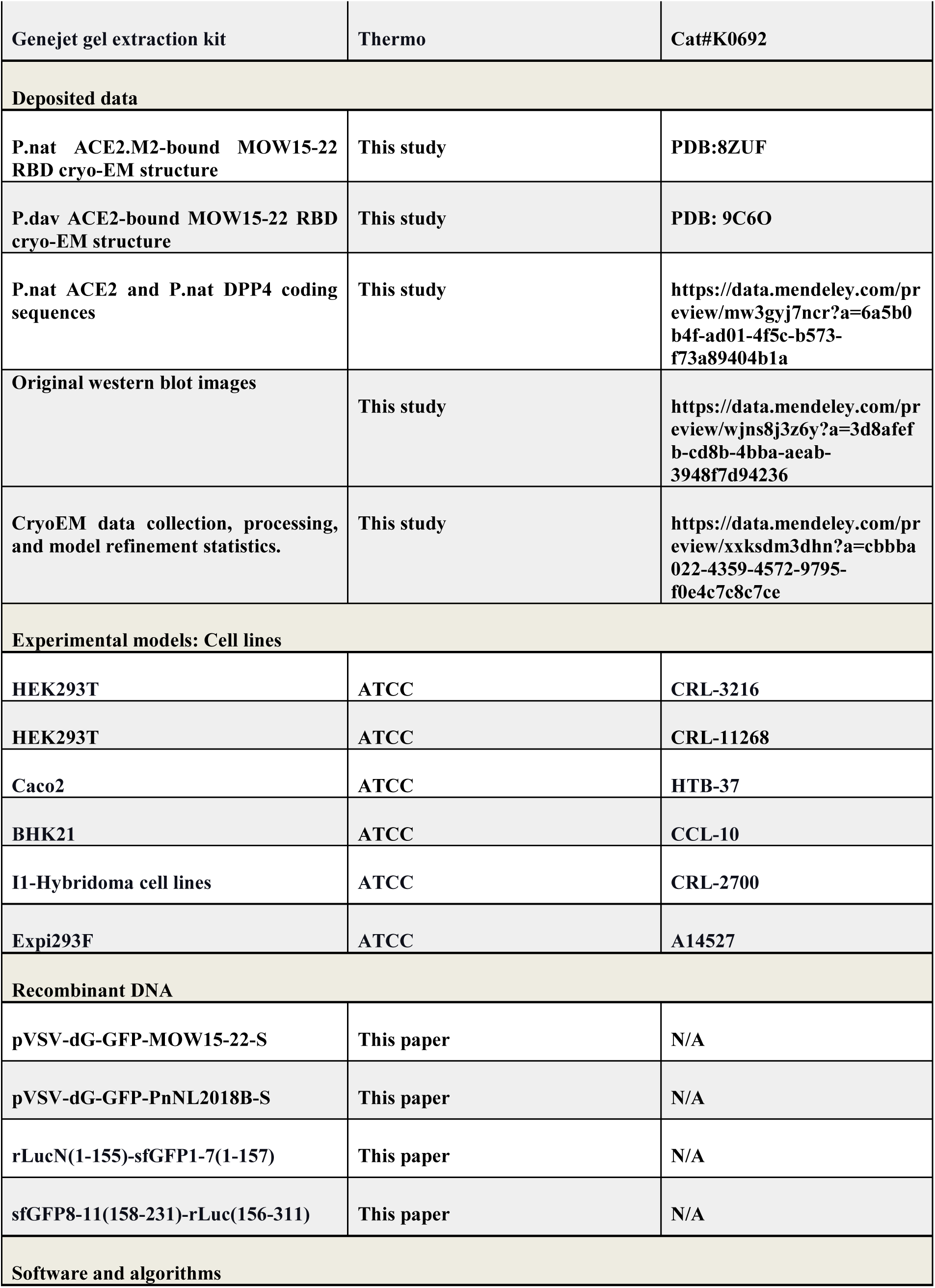
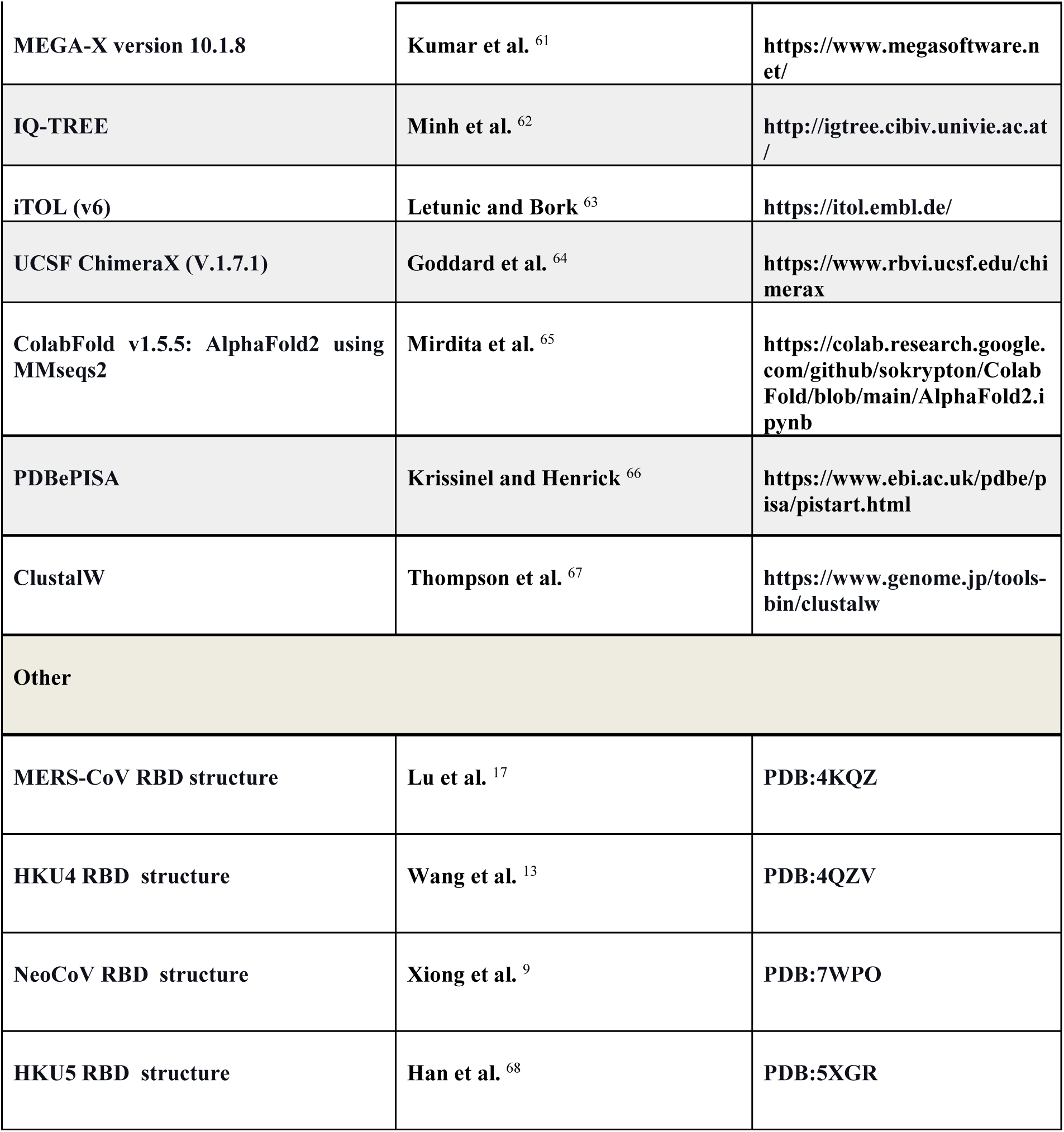

## RESOURCE AVAILABILITY

### Lead contact

Further information and requests for resources and reagents should be directed to and will be fulfilled by the lead contact, Huan Yan (严欢)

### Materials availability

All plasmids generated in this study are available with a completed Materials Transfer Agreement. Correspondence and requests for materials can be addressed to the lead contact.

### Data and code availability

- ● The cryoEM structures have been deposited to the electron microscopy data bank and protein data bank with accession numbers EMD-45253, PDB-9C6O (P.dav ACE2-bound MOW15-22 RBD) and EMD-60483, PDB-8ZUF (P.nat ACE2.M2-bound MOW15-22

RBD). Original western blot images, cryo-EM data statistics, P.nat ACE2 and P.nat DPP4 coding sequences have been deposited at Mendeley and are publicly available as of the date of publication. The DOI is listed in the key resources table. Accession numbers are listed in the key resources table. Microscopy data reported in this paper will be shared by the lead contact upon request.

● This paper does not report original code.
● Any additional information required to reanalyze the data reported in this paper is available from the lead contact upon request.

## EXPERIMENTAL MODEL AND STUDY PARTICIPANT DETAILS

### Cell lines and culture conditions

HEK293T (CRL-3216), HEK293T (ATCC, CRL-11268) Caco2 (HTB-37), BHK21 (CCL-10), and I1-Hybridoma (CRL-2700) cell lines were obtained from the American Type Culture Collection (ATCC). Expi293F (A14527) cells were obtained from Thermo Fisher Scientific. These cells were maintained in Dulbecco’s Modified Eagle Medium (DMEM, Monad, China) supplemented with 1% PS (Penicillin/Streptomycin) and 10% Fetal Bovine Serum (FBS). The I1-Hybridoma cell line, which produces a neutralizing antibody targeting the VSV glycoprotein (VSV-G), was cultured in Minimum Essential Medium (MEM) with Earles’s balances salts, 2.0 mM of L-glutamine (Gibico), and 10% FBS. All cell lines were cultured at 37℃ with 5% CO2 and routinely passaged every 2-3 days. HEK293T or Caco2 stable cell lines overexpressing various receptors were generated using lentivirus transduction and antibiotic selection. The stable cells based on HEK293T or Caco2 were selected and maintained in the growth medium with puromycin (1 μg /ml).

## METHOD DETAILS

### Plasmids and vectors

Plasmids expressing wild-type (WT) or mutated bat and non-bat mammalian ACE2 orthologs ^24,33^ were constructed by inserting human codon-optimized sequences with/without specific mutations into a lentiviral transfer vector (pLVX-EF1a-Puro, Genewiz) with C-terminus 3×FLAG tags (DYKDHD-G-DYKDHD-I-DYKDDDDK) and single FLAG tags (DYKDDDDK) for non-bat mammalian ACE2 orthologs^24^. P.nat ACE2 and DPP4 coding sequences are retrieved from the genome of *Pipistrellus nathusii* (GCA_963693515.1), P.nat ACE2 and P.nat DPP4 coding sequences have been deposited at Mendeley with DOI listed in the key resources table. ^31^. For pseudovirus production, human codon-optimized spike sequences of MOW15-22 (USL83011.1), PnNL2018B (WDE20340.1), SARS-CoV-2 (YP_009724390.1) carrying D614G mutation, MERS-CoV (YP_009047204.1), HKU4 (AWH65899), NeoCoV (AGY29650.2) and HKU31 (QGA70692.1) were cloned into the pCAGGS vector with C-terminal deletions (residues 13-15) for improving the pseudovirus assembly efficiency. The DNA fragments for cloning ACE2 chimera or mutants were generated by overlap extension PCR or gene synthesis and verified by commercial DNA sequencing. For the expression of recombinant CoVs RBD- hFc fusion proteins, plasmids were constructed by inserting NeoCoV RBD (residues 380-585), MOW15-22 RBD (residues 360-610aa), PnNL2018B RBD (residues 361-606) coding sequences into the pCAGGS vector containing an N-terminal CD5 secretion signal peptide (MPMGSLQPLATLYLLGMLVASVL) and C-terminal hFc-twin-strep tandem tags for purification and detection. Plasmids expressing soluble ACE2 ectodomain proteins were generated by inserting sequences from hACE2 (residues 18-740), P.dav ACE2 (residues 18- 738), and P.nat ACE2 (residues 18-739) into pCAGGS vector, with an N-terminal CD5 secretion signal peptide and a C-terminal twin-strep-3×FLAG tag (WSHPQFEKGGGSGGGSGGSAWSHPQFEK- GGGRSDYKDHDGDYKDHDIDYKDDDDK)^9^. The construct for expressing soluble P.nat ACE2.M1 was generated by overlapping PCR to replace residues of 404-430 with P.dav ACE2 corresponding sequences. The P.nat ACE2.M2 was generated by further introducing a Q598L point mutation to P.nat ACE2.M1. For CryoEM analysis, the P.dav ACE2 ectodomain (residues 17-723) construct harbors an N-terminal signal peptide (MPMGSLQPLATLYLLGMLVASVL) with avi tag followed by seven residues flexible linker and C-terminal octa-histidine tag was codon-optimized, synthesized, and inserted the pcDNA3.1(+) vector by Genscript. The MOW15- 22 RBD construct encoding S residues 351-599 containing N-terminal signal peptide (MGILPSPGMPALLSLVSLLSVLLMGCVA), with an avi tag followed by eight residues flexible linker and C-terminal octa-histidine tag were codon optimized, synthesized, and inserted into the pcDNA3.1(+) vector by Genscript.

### Protein expression and purification

HEK293T cells were transfected with corresponding plasmids using GeneTwin reagent (Biomed, TG101-01). Subsequently, the culture medium of the transfected cells was replenished with the SMM 293-TII Expression Medium (Sino Biological, M293TII) 4-6 hours post-transfection, and the protein-containing supernatant was collected every three days for 2-3 batches. Antibodies and recombinant RBD-hFc proteins were purified using Pierce Protein A/G Plus Agarose (Thermo Scientific, 20424). In general, Fc-containing proteins were enriched by the agarose, washed with wash buffer (100 mM Tris/HCl, pH 8.0, 150 mM NaCl, 1 mM EDTA), eluted using the Glycine buffer (100 mM in H2O, pH 3.0), and immediately neutralized with 1/10 volume of 1M Tris-HCI, pH 8.0 (15568025, Thermo Scientific). Proteins with twin-strep tag were purified using Strep-Tactin XT 4Flow high-capacity resin (IBA, 2-5030-002), washed by wash buffer (100 mM Tris/HCl, pH 8.0, 150 mM NaCl, 1 mM EDTA), and then eluted with buffer BXT (100 mM Tris/HCl, pH 8.0, 150 mM NaCl, 1 mM EDTA, 50 mM biotin). All eluted proteins were concentrated using Ultrafiltration tubes, buffer-changed to PBS, and stored at -80℃. Protein concentrations were determined by the Omni-Easy Instant BCA Protein Assay Kit (Epizyme, ZJ102).

### Antigen-hFc live-cell binding assay

RBD- or S1-hFc live-cell binding assays were conducted following a previously described protocol^33^. The coronavirus RBD- or S1-hFc recombinant proteins were diluted in DMEM at indicated concentrations and incubated with HEK293T cells transiently expressing different ACE2 for 30 minutes at 37℃ at 36 hours post-transfection. Subsequently, cells were washed once with Hanks’ Balanced Salt Solution (HBSS) and incubated with 1 μg/mL of Alexa Fluor 488-conjugated goat anti-human IgG (Thermo Fisher Scientific; A11013) diluted in HBSS/1% BSA for 1 hour at 37℃. After another round of washing with HBSS, the cell nuclei were stained with Hoechst 33342 (1:10,000 dilution in HBSS) for 30 minutes at 37℃. The images were captured using a fluorescence microscope (MI52-N). The relative fluorescence intensities (RFUs) of the stained cells were determined by a Varioskan LUX Multi-well Luminometer (Thermo Scientific). The heatmap presentation in Figure 2A, B, and S3C, D, was set based on RFUs subjected with background RLUs in cells without ACE2 expression. Color ranges and thresholds were adjusted to show binding efficiencies consistent with the fluorescence images captured by the microscope.

### Flow cytometry

To analyze S1-hFc binding through flow cytometry, HEK293T cells transiently expressing the indicated ACE2 orthologs were detached with 5 mM EDTA/PBS at 36 hours post-transfection. The cells were washed twice with cold PBS and incubated with MOW15-22 or PnNL2018B S1- hFc proteins at 2 μg/mL concentrations at 4℃ for 30 minutes. Subsequently, cells were incubated with Alexa Fluor 488-conjugated goat anti-human IgG to stain the RBD (Thermo Fisher Scientific; A11013) at 4℃ for 1 hour. Afterward, cells were fixed with 4% PFA, permeabilized with 0.25% Triton X-100, blocked with 1% BSA/PBS at 4℃, and then incubated with mouse antibody M2 (Sigma-Aldrich, F1804) diluted in PBS/1% BSA for 1 hour at 4℃, followed by incubation with Alexa Fluor 594-conjugated goat anti-mouse IgG (Thermo Fisher Scientific; A32728) diluted in 1% BSA/PBS for 1 hour at 4℃. For all samples, 10,000 ACE2- expressing live cells (gated based on SSC/FSC and FLAG-fluorescence intensity and SSC/FSC) were analyzed using a CytoFLEX Flow Cytometer (Beckman).

### Biolayer interferometry (BLI) binding assay

Protein binding kinetics was assessed using BLI assays with the Octet RED96 instrument (Molecular Devices) at 25℃, and shaking at 1000 rpm. Specifically, RBD-hFc or S1-hFc recombinant proteins were diluted to 20 μg/mL and immobilized on Protein A (ProA) biosensors (ForteBio, 18-5010), which were then incubated with soluble dimeric bat ACE2-ectodomain proteins, which were two-fold serial-diluted in kinetic buffer (PBST), starting from 1,000 nM. A well incubated with kinetic buffer (PBST) only serves as a background control. The kinetic parameters and the apparent binding affinities (due to ACE2 dimerization) between the RBD-hFc and ACE2 were analyzed using Octet Data Analysis software 12.2.0.20 with global curve fitting using a 1:1 binding model.

P.dav ACE2 binding to the MOW15-22 RBD (Figure 2C) was performed at 30°C and shaking at 1,000 rpm. The biotinylated MOW-15-22 RBD was diluted to 10µg/mL in 10× Octet kinetics buffer (Sartorius) and loaded onto hydrated Streptavidin biosensors to 1nm shift, equilibrated in 10x Octet kinetics buffer for 60 seconds, and dipped into P. davyi ACE2 at 600nM, 200nM, 66.6nM and 22nM for 300s to observe association. Dissociation was followed by dipping biosensors in a 10× Octet kinetics buffer for 300s. Baseline subtraction was done by subtracting the response from unloaded SA tips dipping into 600nM P. davyi ACE2. In addition, association phases were aligned to 0 seconds and 0 response in Octet Data Analysis HT software and the processed results were exported. Global fitting using a 1:1 binding model was carried out using the Octet Data Analysis HT software Sensorgrams and plotted in GraphPad Prism10.

### Cell-cell fusion assays

A cell-cell fusion assay based on dual-split proteins (DSPs) was conducted in Caco2 cells stably expressing ACE2 receptors. To assess the S glycoprotein-receptor interaction-mediated membrane fusion between cells, group A cells were transfected with S glycoprotein and rLucN(1-155)-sfGFP1-7(1-157) expressing plasmids, while group B cells were transfected with S glycoprotein and sfGFP8-11(158-231)-rLuc(156-311) expressing plasmids. After 12 hours of transfection, both groups of cells were trypsinized, mixed, and seeded into a 96-well plate at 8 × 10^4^ cells per well. Subsequently, the cells were washed once with DMEM and then incubated with DMEM with or without indicated concentrations of TPCK-treated trypsin (Sigma-Aldrich, T8802) for 10 min at room temperature at 24 h post-transfection. Six hours later, nuclei were stained with Hoechst 33342 (1:5,000 dilution in HBSS) for 30 min at 37 °C, and images of syncytia formation with green fluorescence were subsequently captured using a fluorescence microscope (MI52-N; Mshot). To measure live-cell luciferase activity after cell-cell fusion, 20 μM of EnduRen live-cell substrate (Promega, E6481) was added to the cells in DMEM and incubated for at least 1 hour before detection using the Varioskan LUX Multi-well Luminometer (Thermo Fisher Scientific).

### Pseudovirus production

VSV-dG-based pseudovirus (PSV) carrying trans-complementary S glycoproteins from various coronaviruses were produced following a modified protocol as previously described^69^. Briefly, HEK293T cells were transfected with plasmids expressing coronaviruses S glycoproteins. At 24 hours post-transfection, cells were transduced with 1.5×10^6^ TCID50 VSV-G glycoprotein- deficient VSV expressing GFP and firefly luciferase (VSV-dG-GFP-fLuc, constructed and produced in-house) diluted in DMEM with 8 μg/mL polybrene for 4-6 hours at 37 ℃. After three PBS washes, the culture medium was replenished with either DMEM+10% FBS or SMM 293- TII Expression Medium (Sino Biological, M293TII), along with the presence of the neutralizing antibody (from I1-mouse hybridoma) targeting the VSV-G to eliminate the background due to any remaining VSV-dG-GFP-fLuc. Twenty-four hours later, the pseudovirus containing supernatant was clarified through centrifugation at 12,000 rpm for 5 minutes at 4℃, aliquoted, and stored at -80℃. The TCID50 of the PSV was calculated using the Reed-Muench method^70,71^. Heatmap presentation in Figure 3A, 3B, and S3A was set based on RLU with thresholds set based on entry background in cells without ACE2 expression. The value of P.nat ACE2 was set as a baseline of 100%, and the color of the largest value was set based on the ortholog showing the highest RLU, unless otherwise specified.

### Single-round pseudovirus entry assay

Single-round pseudovirus entry assays were conducted using cells transiently or stably expressing different ACE2 orthologs. Approximately 3×10^4^ trypsinized cells were incubated with pseudovirus (2×10^5^ TCID50/100 μL) in a 96-well plate to facilitate attachment and viral entry simultaneously. Before inoculation, pseudoviruses were typically treated with 100 μg/mL TPCK-trypsin (Sigma-Aldrich, T8802). Specifically, pseudoviruses produced in serum-free SMM 293-TII Expression Medium were incubated with TPCK-treated trypsin for 10 minutes at room temperature, and the proteolytic activity was neutralized by FBS in the culture medium. Intracellular luciferase activity (Relative light units, RLU) was measured using the Bright-Glo Luciferase Assay Kit (Promega, E2620) and detected with a GloMax 20/20 Luminometer (Promega) or Varioskan LUX Multi-well Luminometer (Thermo Fisher Scientific) at 18 hours post-infection.

### pcVSV-CoV amplification assay

Plasmids for rescuing propagation-competent (pc) VSV-CoV expressing MOW15-22 and PnNL2018B S glycoproteins (pVSV-dG-GFP-MOW15-22-S and pVSV-dG-GFP-PnNL2018B-S) were generated by replacing the fLuc coding sequences with coronavirus spike coding sequences based on the vector pVSV-dG-GFP-fLuc. The propagation-competent recombinant VSVs were created by replacing the VSV-G gene with genomically encoded S glycoprotein genes and additionally incorporating a GFP-expressing cassette for visualization. Reverse genetics was applied to rescue pcVSV pseudotypes expressing MOW15-22 and PnNL2018B S glycoproteins along with a GFP reporter, following a modified protocol from previous descriptions^69^. Specifically, BHK21 cells, seeded in a 6-well plate at 80% confluence, were inoculated with 5 MOI of recombinant vaccinia virus expressing T7 RNA polymerase (vvT7, a kind gift from Mingzhou Chen’s lab, Wuhan University) for 45 minutes at 37°C. After removing vvT7, cells were subsequently transfected with pVSV-dG-GFP-S vector plasmids and helper plasmids (pVSV-dG-GFP-S: pBS-N: pBS-P: pBS-G: pBS-L=5:3:5:8:1). pcVSV-CoV containing supernatant (P0) was collected 48 hours post-transfection and 0.22-μm filtered to remove vvT7. For enhanced virus amplification, the passage 1 (P1) virus carrying VSV-G proteins was produced by inoculating P0 supernatant with Caco2 cells 24 hours post-transfection of plasmids expressing VSV-G protein. Subsequently, P1 viruses were further amplified in Caco2 cells stably expressing P.dav ACE2 or L.bor ACE2, without the ectopic expression of VSV-G and in the presence of anti-VSVG (I1-Hybridoma supernatant), producing passage 2 (P2) viruses carrying MOW15-22 and PnNL2018B S glycoproteins, respectively. In a typical virus amplification assay, 3×10^4^ trypsinized cells stably expressing the indicated ACE2 were incubated with pcVSV-CoV (1×10^4^ TCID50/100 μL) in a 96-well plate with or without TPCK-treated trypsin treatment. GFP images or RFU were collected at indicated time points post-infection by a fluorescence microscope (MI52-N) or a Varioskan LUX Multi-well Luminometer (Thermo Fisher Scientific).

### Pseudotyped virus neutralization assays

For soluble ACE2 neutralization assays, serial dilutions of recombinant proteins were prepared in DMEM. Pseudotyped viruses (1 × 10^5^ TCID50 per well) were mixed with 25 µl of each dilution for 1 hour at 37 °C, and then incubated P.nat ACE2-expressing Caco2 cells seeded at 2 × 10^4^ cells per well in a 96-well plate. After 16–20 hours post-infection, the luciferase activity was measured as described in the single-round pseudotype virus entry assays. For S2P6, 76E1 antibody neutralization assays, pseudoviruses were incubated with 3-fold serial dilutions of antibodies in DMEM for 1 hour at 37 °C, and then applied to HEK293T stably expressing the P.dav or L.bor ACE2. For the H11B11 neutralization assay, Caco2 cells stably expressing the hACE2.3M were seeded on the 96-well plate one day before infection, 3-fold serial dilutions of H11B11 were diluted in the culture medium and then incubated with the cells for 1 hour at 37 °C. Pseudoviruses were then added to the cells with the previous H11B11-containing medium. Neutralizing efficiencies were assessed at 18 hours post-infection by detecting the intracellular luciferase activity (RLU).

### Western blot

For detecting the cellular expression of ACE2 or DPP4 receptors with C-terminal FLAG tags, cells at 24 hours post-transfection were lysed in 1% TritonX/PBS+1 mM PMSF (Beyotime, ST506) for 10 minutes at 4℃. The lysate was clarified after centrifugation at 12,000 rpm for 5 minutes at 4℃ and then incubated at 98℃ for 10 minutes after mixing with the 1/5 volume of 5×SDS loading buffer. Following gel electrophoresis and membrane transfer, the membranes were blocked with 5% skimmed milk in PBST for 2 hours at room temperature. Subsequently, the membrane was incubated with 1 μg/mL anti-FLAG mAb (Sigma, F1804) or anti-β-tubulin (Immmuno Way, YM3030) mAb diluted in PBST containing 1% milk overnight at 4℃. After four washes with PBST, the blots were incubated with Horseradish peroxidase (HRP)-conjugated secondary antibody AffiniPure Goat Anti-Mouse in 1% skim milk diluted in PBST and incubated for one hour at room temperature. Finally, the blots were washed four times again by PBST and visualized using an Omni-ECL Femto Light Chemiluminescence Kit (EpiZyme, SQ201) through a ChemiDoc MP Imaging System (Bio-Rad). To examine the S glycoprotein packaging efficiency, the PSV-containing supernatant was concentrated using a 30% sucrose cushion (30% sucrose, 15 mM Tris-HCl, 100 mM NaCl, 0.5 mM EDTA) at 20,000×g for 30 minutes at 4℃. The concentrated virus pellet was re-suspended in 1×SDS loading buffer and incubated at 95℃ for 10 minutes, followed by western blot detecting the S glycoproteins by C- terminal HA tags and with the VSV-M serving as a loading control. Original western blot images have been deposited at Mendeley with DOI listed in the key resources table.

### Immunofluorescence assay

Immunofluorescence assays were conducted to determine the expression levels of ACE2 orthologs with C-terminal fused FLAG tags. Specifically, the transfected cells were fixed and permeabilized by incubation with 100% methanol for 10 minutes at room temperature. Subsequently, the cells were incubated with a mouse antibody M2 (Sigma-Aldrich, F1804) diluted in PBS/1% BSA for one hour at 37℃. After a PBS wash, the cells were incubated with Alexa Fluor 594-conjugated goat anti-mouse IgG (Thermo Fisher Scientific, A32742) secondary antibody diluted in 1% BSA/PBS for one hour at 37℃. The images were captured and merged with a fluorescence microscope (Mshot, MI52-N) after the nucleus was stained blue with Hoechst 33342 reagent (1:5,000 dilution in PBS).

### Recombinant glycoprotein production for cryo-EM analysis

The P.dav ACE2 and P.nat ACE2.M2 ectodomain or MOW15-22 RBD were produced and purified using Expi293F or ExpiCHO-S cells. Expi293F cells were grown to a density of 3 × 10^6^ cells/mL and transfected using the ExpiFectamine 293 Transfection Kit (ThermoFisher Scientific) and expression was carried out for 4 days post-transfection at 37°C with 8% CO2. The P.dav ACE2 ectodomain and MOW15-22 RBD were purified from clarified supernatants using HisTrap HP affinity columns (Cytiva) and washed with ten column volumes of 10 mM imidazole, 25 mM sodium phosphate pH 8.0, and 300 mM NaCl before elution with two column volumes of 300 mM imidazole, 25 mM sodium phosphate pH 8.0, and 300 mM NaCl. Purified P.dav ACE2 ectodomain or MOW15-22 RBD were then further purified by size exclusion chromatography using a Superdex 200 Increase 10/300 GL column (Cytiva) and concentrated using centrifugal filters (Amicon Ultra) before being flash frozen. P.nat ACE2.M2 with twin-strep tag was purified with Strep-Tactin XT 4Flow high-capacity resin (IBA, 2-5030-002), washed with ten column volumes of wash buffer (100 mM Tris/HCl, pH 8.0, 150 mM NaCl), and then eluted with a buffer containing 100 mM Tris/HCl, pH 8.0, 150 mM NaCl, 50 mM biotin. P.nat ACE2.M2 and MOW15-22 RBD were initially mixed at a molar ratio of 1:1.5 and incubated for 1h on ice, and then were further purified in a Superose 6 Increase 10/300 GL column (Cytiva) with a buffer containing 20 Tris/HCl, pH 8.0, 150 mM NaCl. The fractions containing the complex were collected and concentrated to 1 mg/ml for further use.

### CryoEM sample preparation, data collection, and data processing

Complex formation was performed by mixing a 1:2.5 molar ratio of the P.dav ACE2 ectodomain (residues 17-723 with C-terminal octa-histidine tag) with the MOW15-22 RBD (residues 351- 599 with C-terminal octa-histidine tag) before incubation for 1 hour at room temperature. CryoEM grids of the complex were prepared using two separate methods and data were combined during data processing. For the first dataset, 3 µL of 4 mg/ml complex with 6 mM 3- [(3-Cholamidopropyl)dimethylammonio]-2-hydroxy-1-propanesulfonate (CHAPSO) were applied onto freshly glow discharged R 2/2 UltrAuFoil grids^72^ prior to plunge freezing using a vitrobot MarkIV (ThermoFisher Scientific) with a blot force of 0 and 5.5 sec blot time at 100% humidity and 22°C. 14,561 movies were collected from UltrAuFoil grids with CHAPSO detergent with a defocus range comprised between -0.2 and -3.5 μm, and stage tilt angle of 0, 20° and 30°^73^. For the second dataset, 3 µL of 0.2 mg/mL complex was added to the glow discharged side of R 2/2 UltrAuFoil grids and 1µL was added to the back side before plunging into liquid ethane using a GP2 (Leica) with 6 sec blot time. 8,238 movies were collected with a defocus range comprised between -0.2 and -3.5 μm. The data were acquired using an FEI Titan Krios transmission electron microscope operated at 300kV and equipped with a Gatan K3 direct detector and Gatan Quantum GIF energy filter, operated in zero-loss mode with a slit width of 20 eV. Automated data collection was carried out using SerialEM^74^ or Leginon^75^ at a nominal magnification of 105,000× with a pixel size of 0.843 Å. The dose rate was adjusted to 9 counts/pixel/s, and each movie was acquired in counting mode fractionated in 100 frames of 40 ms. Movie frame alignment, estimation of the microscope contrast-transfer function parameters, particle picking, and extraction were carried out using Warp^76^. Particles were extracted with a box size of 192 pixels with a pixel size of 1.686Å. Two rounds of reference-free 2D classification were performed using CryoSPARC^77^ to select well-defined particle images from each dataset. Particles belonging to classes with the best resolved RBD and ACE2 density were selected. To improve particle picking further, we trained the Topaz^78^ picker on Warp-picked particle sets belonging to the selected classes after 2D classification. The particles picked using Topaz were extracted and subjected to 2D classification using cryoSPARC, which improved the number of unique 2D views. The two different particle sets picked from Warp and Topaz were merged and duplicate particle picks were removed using a minimum distance cutoff of 160Å. Initial model generation was done using ab-initio reconstruction in cryoSPARC and used as references for a heterogenous 3D refinement in cryoSPARC. After two rounds of ab-initio reconstructions and heterogeneous refinements to remove junk particles, 3D refinement was carried out using non-uniform refinement with per-particle defocus refinement in cryoSPARC^79^ and the particles were transferred from cryoSPARC to Relion using pyem (https://github.com/asarnow/pyem) to be subjected to the Bayesian polishing procedure implemented in Relion^80^ during which particles were re-extracted with a box size of 320 pixels and a pixel size of 1.0 Å. After ab-initio reconstructions and heterogeneous refinements to remove RBD unbound class, subsequent 3D refinement used non-uniform refinement along with per-particle defocus refinement in cryoSPARC. To further improve the density of the RBD:ACE2 interface, local refinement was performed using cryoSPARC with a soft mask comprising one ACE2 peptidase domain and the bound RBD. Local resolution estimation, filtering, and sharpening were carried out using cryoSPARC to yield the final reconstruction at 2.8 Å resolution comprising 705,956 particles. Reported resolutions are based on the 0.143 gold- standard Fourier shell correlation (FSC) criterion and Fourier shell correlation curves were corrected for the effects of soft masking by high-resolution noise substitution^81,82^.

For P.nat ACE2.M2 and MOW15-22 RBD complex, 3.5 μL of the purified complex at 0.8 mg/mL was applied to glow-discharged Cu R1.2/1.3 holey carbon grid (200 mesh, Quantifoil) with a final concentration of 1% glycerol. After incubation for 20 s, the grids were blotted with force 0 for 2 s at 4°C and 100% humidity, and plunge-frozen into liquid ethane using Vitrobot Mark IV (FEI Thermo Fisher). Grids were then transferred to a CRYO ARM 300 electron microscope (JEOL, Japan) operating at 300 kV equipped with a K3 direct electron detector (Gatan, USA). Cryo-EM images were recorded automatically using Serial-EM software with a super-resolution pixel size of 0.475 Å /pixel at defocus values ranging from -0.5 to -2.5 μm at a calibrated magnification of 50,000. Data were collected at a frame rate of 40 frames per second. The total electron dose was 40 e-/Å2. 4,214 movies were collected and subjected to patch motion correction and patch CTF estimation in cryoSPARC. Particles were automatically picked with a diameter of 120 Å from 200 micrographs. ∼50,000 particles representing good classes were used to train a topaz model for auto-picking. A subset of 785,038 particles was selected after two rounds of 2D classification, followed by an ab initio reconstitution and a heterogeneous refinement. One class representing intact P.nat ACE2.M2-MOW15-22 RBD complex was further selected for heterogeneous refinement requesting three classes. One class containing 244,733 particles with good features was subjected to non-uniform refinement, yielding a 3.3 Å reconstruction.

### Model building and refinement

UCSF Chimera^83^ was used to rigid-body dock models into the sharpened cryoEM map and adjustments and refinement were carried out with Coot^84^ and Rosetta^85,86^ using sharpened and unsharpened maps. Validation used Molprobity^87^, Phenix^88^ and Privateer^89^.

### Bioinformatic and structural analysis

Sequence alignments of different bats ACE2 were performed using either the MUSCLE algorithm by MEGA-X (version 10.1.8) or ClustalW software (https://www.genome.jp/tools-bin/clustalw) with slight position adjustment to align indels. Phylogenetic trees were generated using the maximal likelihood method in IQ-TREE (http://igtree.cibiv.univie.ac.at/) (1000 Bootstraps) and refined with iTOL (v6) (https://itol.embl.de/). The structures of PnNL2018B, and HKU31 RBDs were predicted using alphaFold2.ipynb-Colaboratory^26^. The structure of NeoCoV RBD & P.pip ACE2 complex (7WPO), MERS-CoV RBD (4KQZ), HKU4 RBD (4QZV), HKU5 RBD (5XGR), MOW15-22 RBD, HKU31 RBD, and PnNL2018B RBD were visualized and analyzed using the ChimeraX(V.1.7.1). Analysis of buried surface area and identification of interface residues were assisted by PISA^66^.

## QUANTIFICATION AND STATISTICAL ANALYSIS

Most experiments related to pseudovirus infection were conducted 2-3 times with 2-4 biological repeats, technical repeats are indicated in the legends. Representative results were shown. All data were presented by MEAN ± SD. Unpaired two-tailed t-tests were conducted for all statistical analyses for two independent groups using GraphPad Prism 8. *P*<0.05 was considered significant. * *p*<0.05, ** *p* <0.01, *** *p* <0.005, and **** *p* <0.001. No data were excluded for data analysis.

---

**Figure S1.**
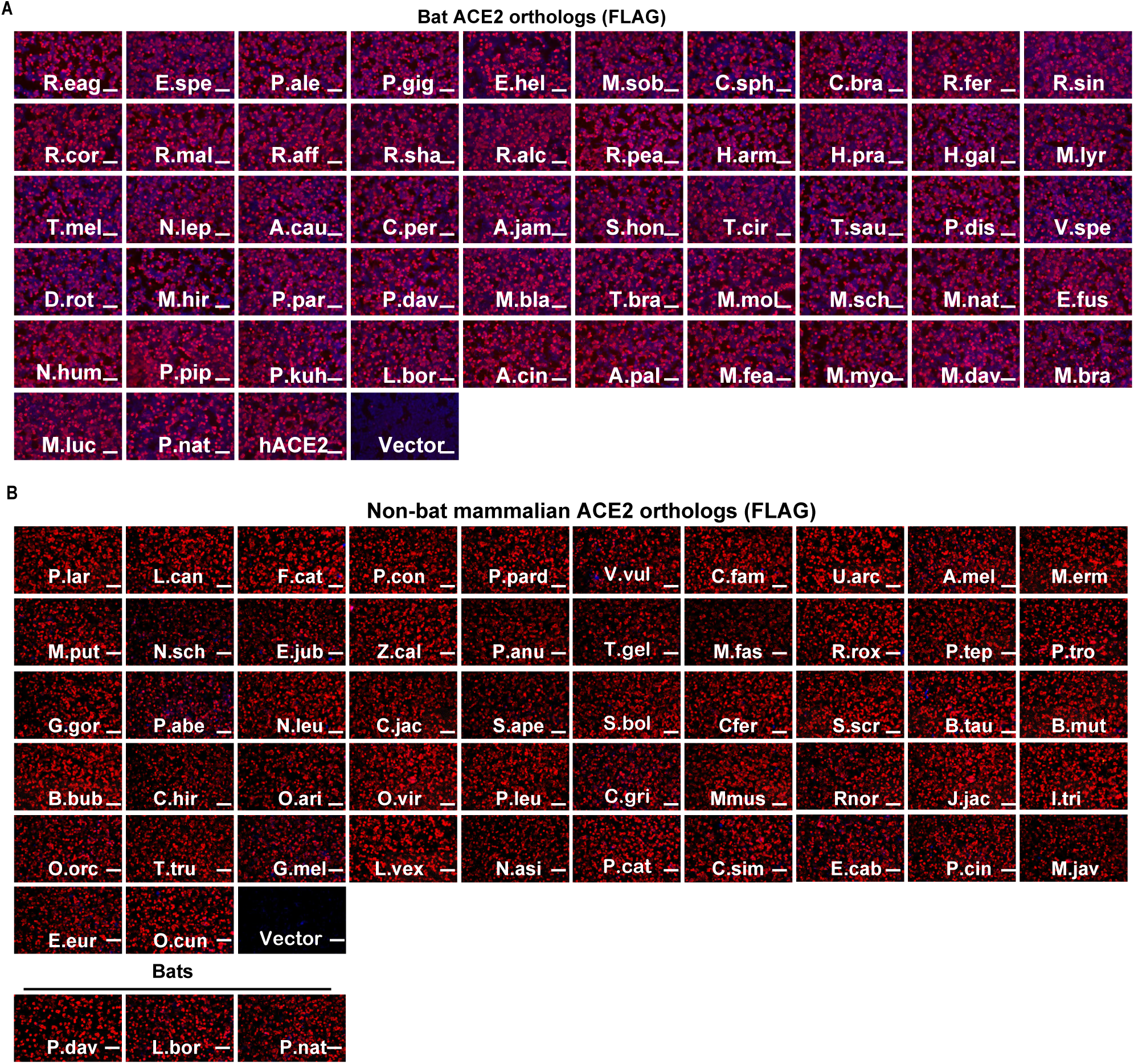
Validation of the expression of ACE2 orthologs from bats or other mammalian species, related to. Figure 3 (**A-B**) Immunofluorescence analyzing the expression of bat (A) or mammalian (B) ACE2 orthologs in HEK293T cells by detecting 3×FLAG tags fused to the C-terminal of the receptors. Scale bars:100 μm.

**Figure S2.**
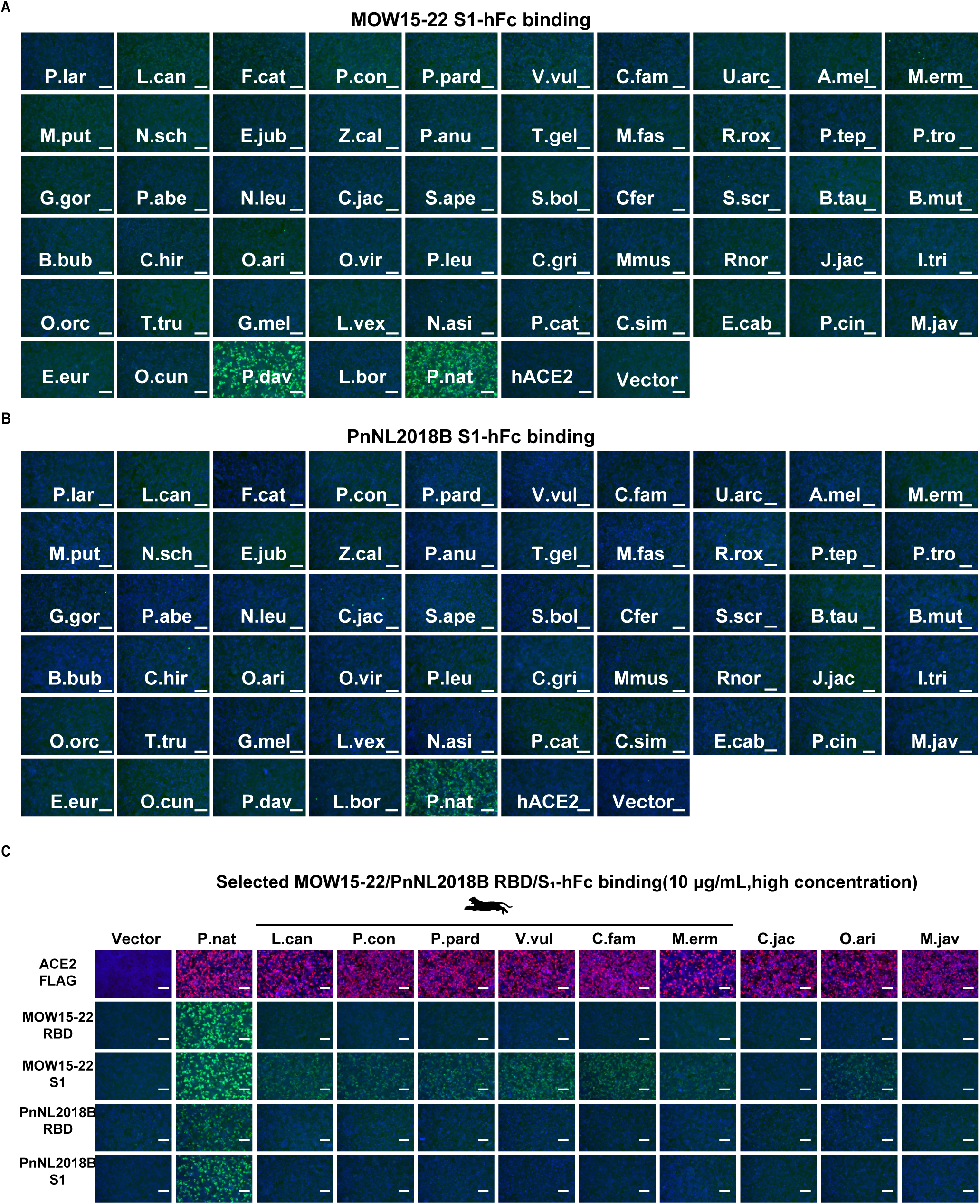
MOW15-22 and PnNL2018B S glycoproteins exhibit narrow ACE2 tropism, related to. Figure 3. (**A-B**) Images showing MOW15-22 (A) and PnNL2018B (B) S1 subunit binding to HEK293T cells transiently expressing the indicated mammalian ACE2 orthologs analyzed by fluorescence microscopy. (**C**) Fluorescence images showing MOW15-22 and PnNL2018B RBD/S1 subunit binding at a higher protein concentration (10 μg/mL) in selected mammalian ACE2 orthologs supporting MOW15-22 entry in Figure. 3B. Scale bars:100 μm.

**Figure S3.**
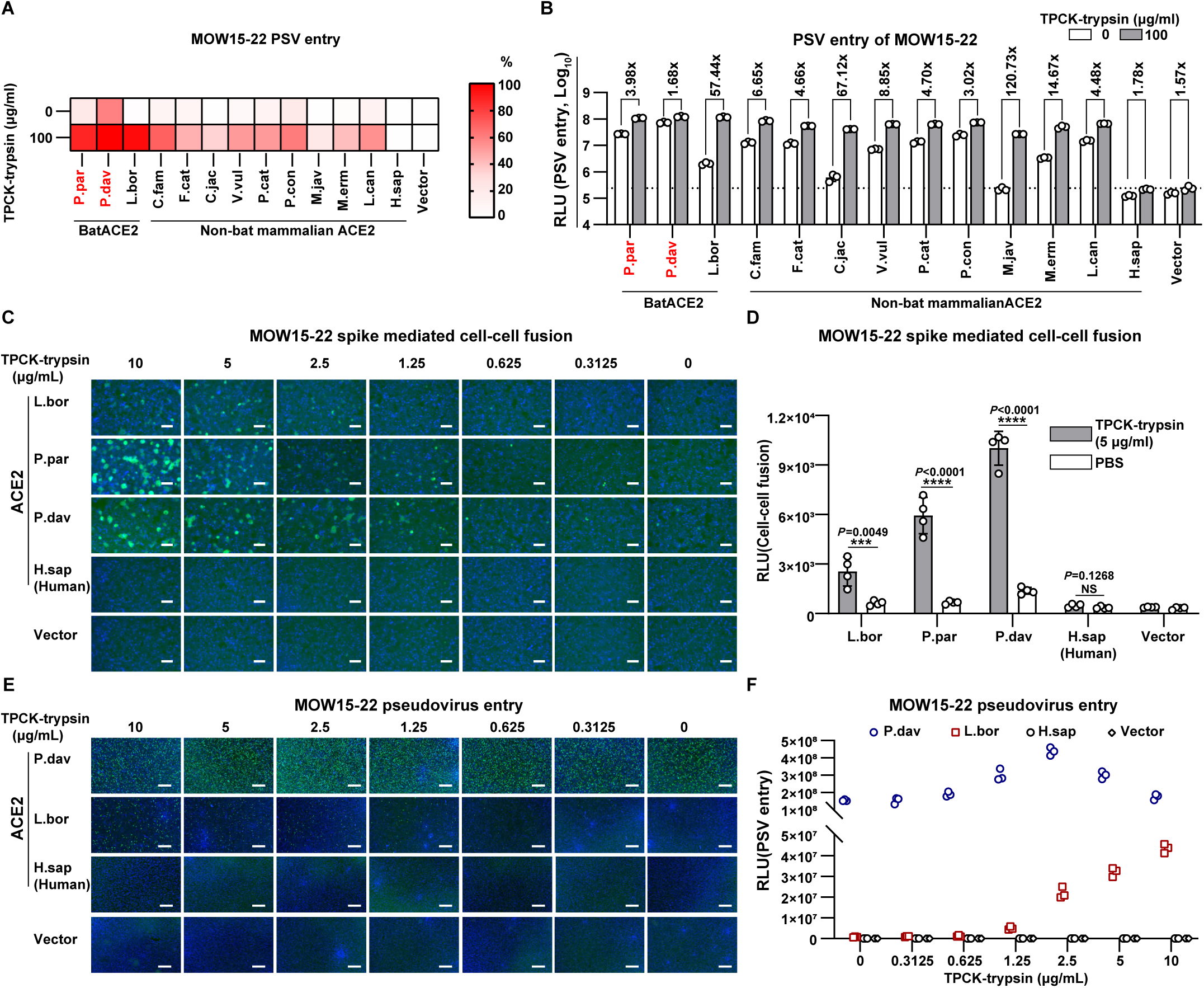
Trypsin-dependence of bat ACE2 mediated MOW15-22 membrane fusion and PSV entry, related to. Figure 3. (**A-B**) Heat map (A) and bar graph (B) of MOW15-22PSV entry efficiency mediated by several ACE2 orthologs in the presence or absence of 100 μg/mL TPCK-treated trypsin. Orthologs showing the highest entry-supporting efficiency (P.dav ACE2 for MOW15-22 and L.bor for PnNL2018B) were set as 100%. Red highlights the two bat ACE2s supporting efficient MOW15-22 RBD binding. The dashed line indicates the background signal. MEAN ± SD and unpaired two-tailed t-tests. n=3 biological replicates. (**C-D**) MOW15-22 spike-mediated cell-cell membrane fusion in HEK293T cells stably expressing the indicated ACE2 orthologs in the presence of various concentrations of TPCK-treated trypsin. Fusion efficiency is indicated by GFP intensity (C) and live-cell Renilla luciferase activity (D) through the reconstitution of dual-split reporter proteins (DSPs). MEAN ± SD and unpaired two-tailed t-tests. n=3 biological replicates. (**E-F**) MOW15-22 PSV entry efficiency in HEK293T cells stably expressing ACE2 orthologs with the indicated concentration of TPCK-treated trypsin, as indicated by GFP intensity (E) and luciferase (F). n=3 biological replicates for F. Scale bars in C and E:200 μm.

**Figure S4.**
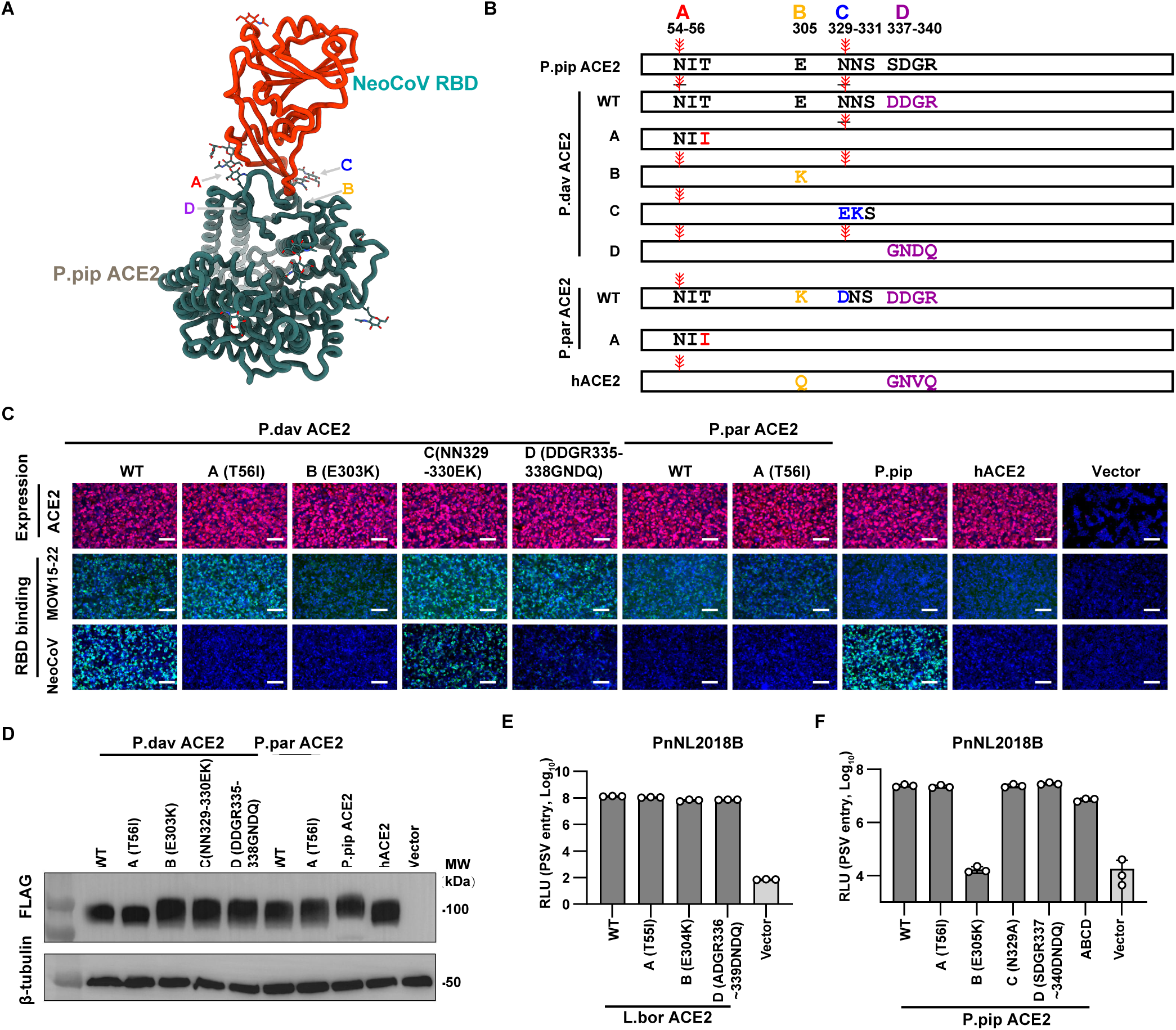
ACE2 determinants critical for NeoCoV/PDF-2180 do not impact MOW15- 22/PnNL2018B ACE2 recognition, related to. Figure 4. (**A**) Structural presentation of the four host range determinants A-D critical for P.Pip ACE2 recognition by NeoCoV (PDB 7WPO). (**B**) Schematic illustration of P.dav and P.par ACE2 mutants with sequences of indicated determinants replaced by residues unfavorable for NeoCoV recognition. Glycosylation sites in determinants A and C are indicated with ￥. (**C**) MOW15-22 and NeoCoV RBD binding to HEK293T cells transiently expressing the indicated wild-type (WT) or mutants ACE2 orthologs. The expression level of indicated ACE2 orthologs was verified by immunofluorescence (C, top panel). Scale bars: 100 μm. (**D**) Western blot analysis of the expression of indicated WT and mutated ACE2 in HEK293T cells. MW: molecular weight. (**E-F**) PSV entry efficiency of PnNL2018B in HEK293T cells transiently expressing the L.bor (E) and P.pip (F) ACE2 mutants affecting NeoCoV receptor recognition. n=3 biological replicates. RLU: relative light unit.

**Figure S5.**
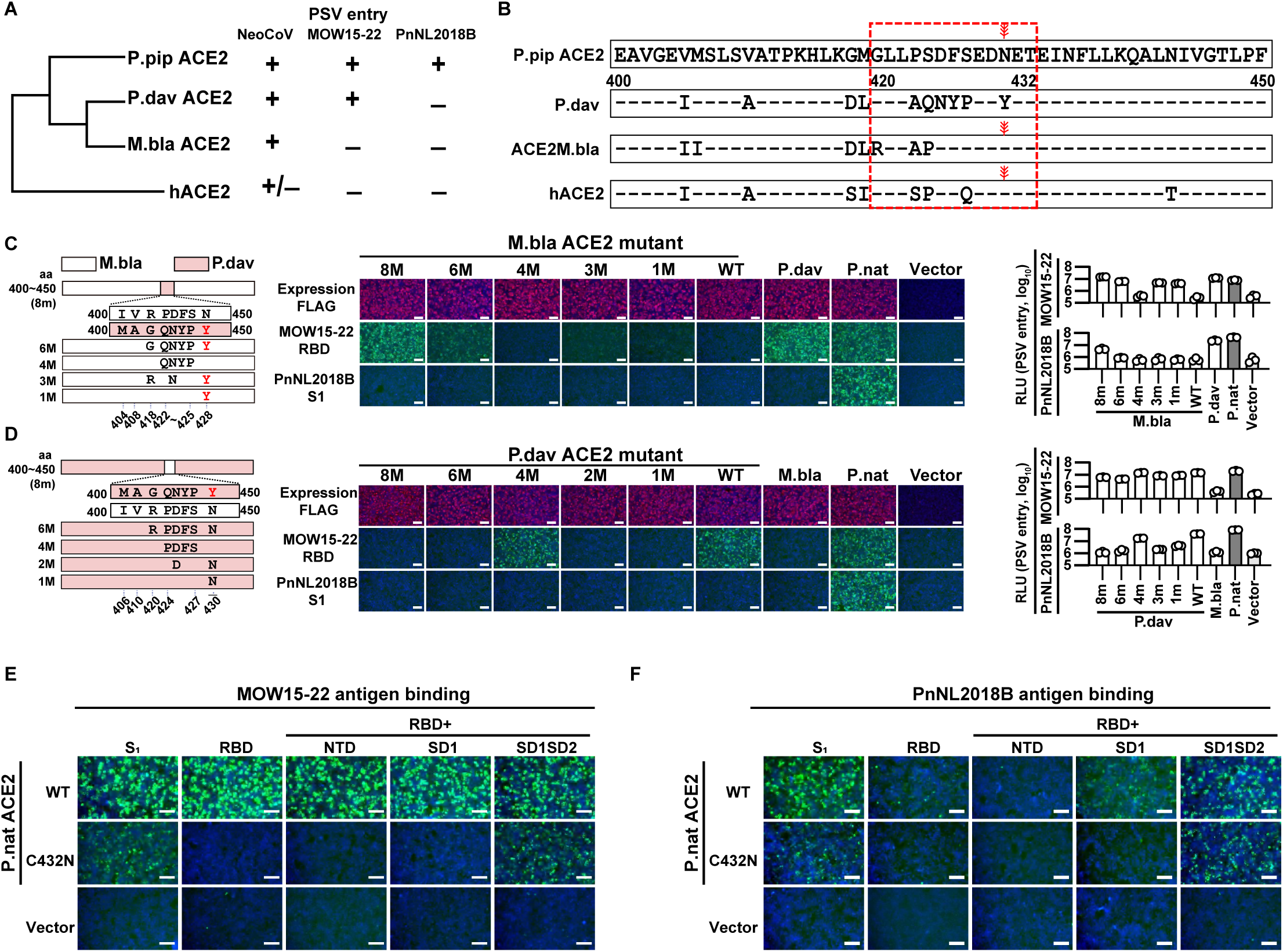
Mapping of ACE2 host range determinants for MOW15-22 and PnNL2018B by genetic swap analysis, related to. Figure 4. (**A-B**) Virus entry-promoting efficiencies (A) and a sequence alignment of residues 400-450 (B) of the selected ACE2 orthologs. Residue numbering is based on P.dav ACE2, and the red dashed box indicates a region participating in host range determination (residues 420-432). ￥: glycosylation, +:supportive, -: not supportive, +/-:weakly supportive. (**C-D**) Determinant mapping based on sequence swaps between residues 400-500 of M.bla and P.dav ACE2. Schematic illustration of chimeric ACE2 mutants enhancing M.bla ACE2 binding (C) abolishing P.dav ACE2 binding (D). RBD binding to (middle) and pseudovirus entry into (right) HEK293T cells transiently expressing the indicated ACE2 swap mutants are shown. MEAN ± SD and unpaired two-tailed t-tests. n=3 biological replicates. (**E-F**) Binding efficiencies of recombinant proteins comprising different domains of the MOW15-22 (C) or PnNL2018B (D) S1 subunit to Caco2 cells stably expressing P.nat ACE2 or a P.nat ACE2 C432N mutant. Scale bars: 100 μm.

**Figure S6.**
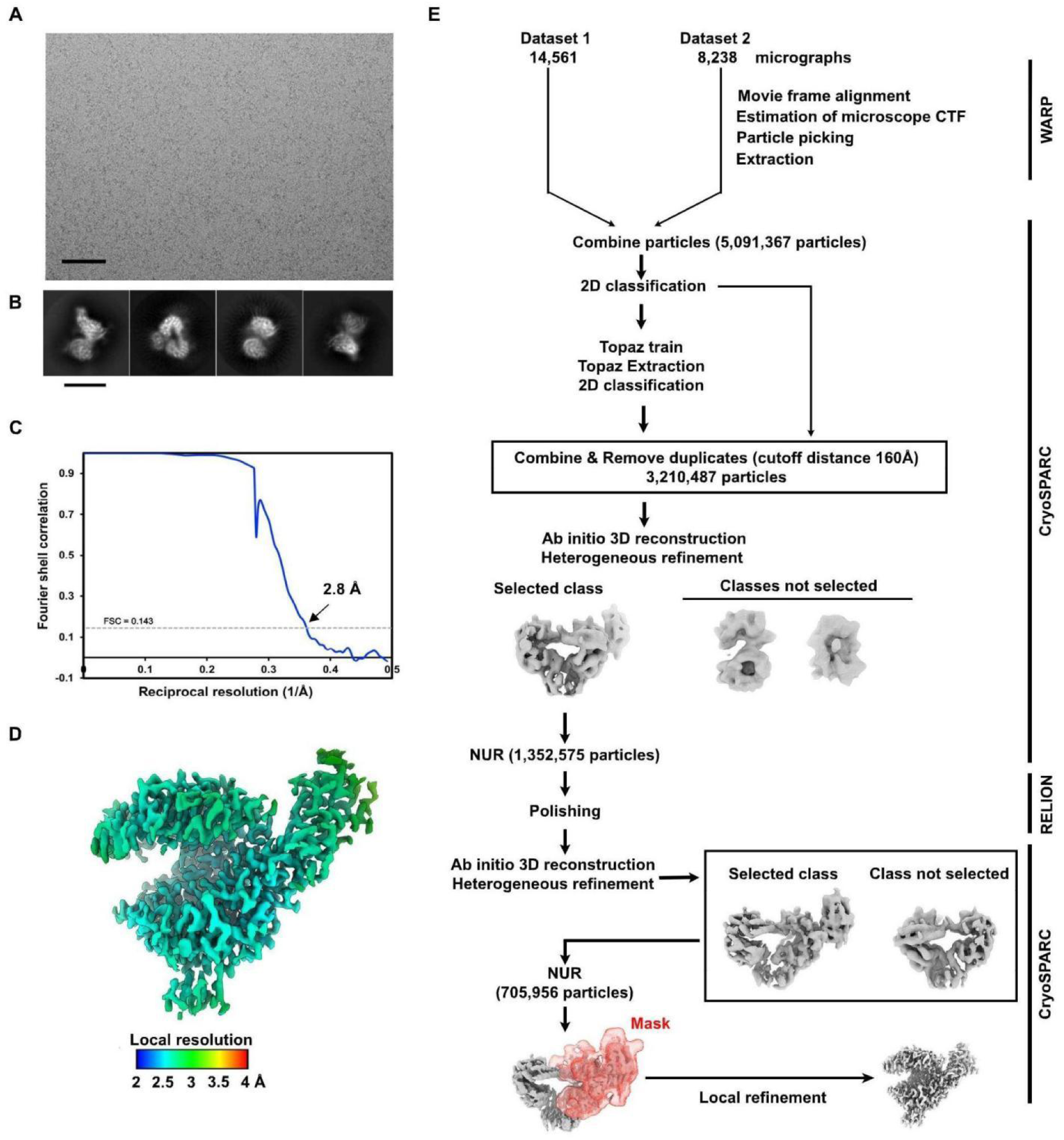
CryoEM data processing workflow for the P.dav ACE2-bound MOW15-22 RBD complex, related to. Figure 5**. (A-B)** Representative electron micrograph (A) and 2D class averages of the P.dav ACE2-bound MOW15-22 RBD complex (B) embedded in vitreous ice. The scale bars represent 100 nm and 200Å, respectively. **(C)** Gold-standard Fourier shell correlation curve. The 0.143 cutoff is indicated by a horizontal dashed line. **(D)** Local resolution estimation was calculated using cryoSPARC and plotted on the sharpened map. **(E)** Data processing flowchart. CTF: contrast transfer function; NUR: non-uniform refinement.

**Figure S7.**
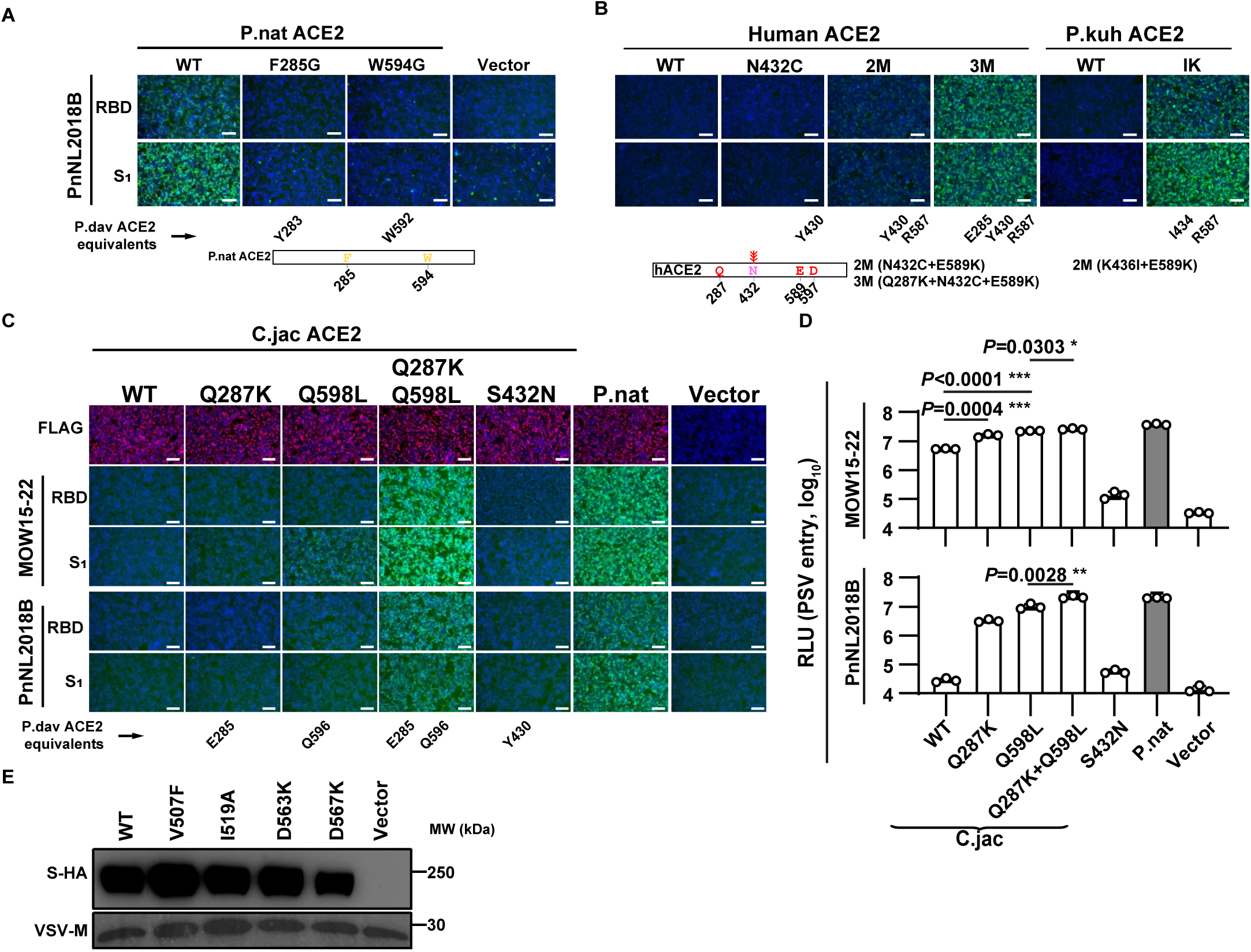
Validation of critical residues for interactions between MOW15-22/PnNL2018B and different ACE2 orthologs, related to. Figure 5**. (A)** P. nat ACE2 mutants with reduced PnNL2018B RBD or S1 binding due to unfavorable substitutions of key interacting residues. (**B**) P.pip, human, and P.kuh ACE2 mutants with enhanced ability to promote PnNL2018B RBD or S1 binding. (**C-D**) C.jac ACE2 mutants promoting enhanced MOW15-22 and PnNL2018B RBD or S1 binding and PSV entry or abolishing receptor function via introducing the N432 glycan knock-in mutation. MEAN ± SD and unpaired two-tailed t-tests. n=3 biological replicates for D. (**E**) VSV packaging efficiencies of MOW15-22 S harboring the indicated mutations.VSV-M serves as a loading control. Scale bars in A, B, and C:100 μm.

**Figure S8.**
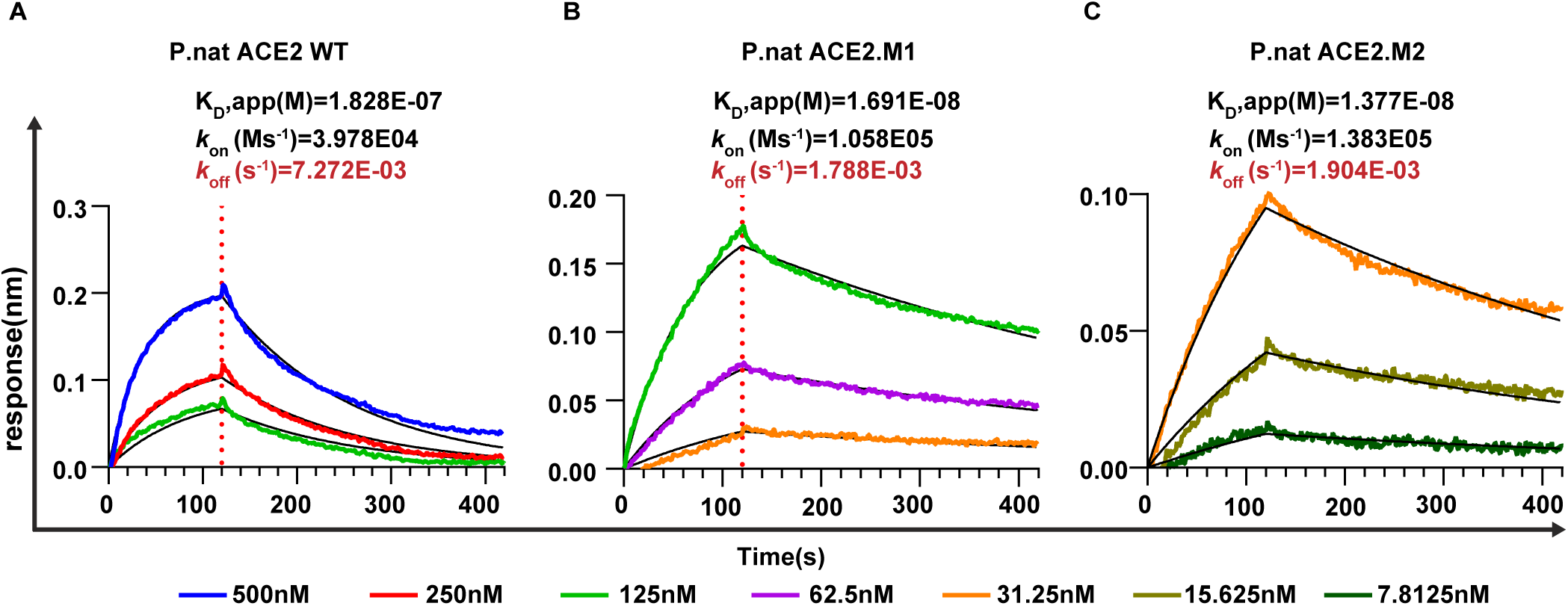
Binding of different soluble P.nat ACE2 mutants to PnNL2018B S1-hFc, related to. Figure 6. BLI analyses of binding kinetics of soluble dimeric ACE2 ectodomains from wildtype (WT) P.nat ACE2 (**A**), P.nat ACE2.M1 (**B**), or P.nat ACE2.M2 (**C**) to the immobilized PnNL2018B S1-hFc. Analysis was conducted with global fitting (1:1 binding model) and the fit to the data is shown in black.

**Figure S9.**
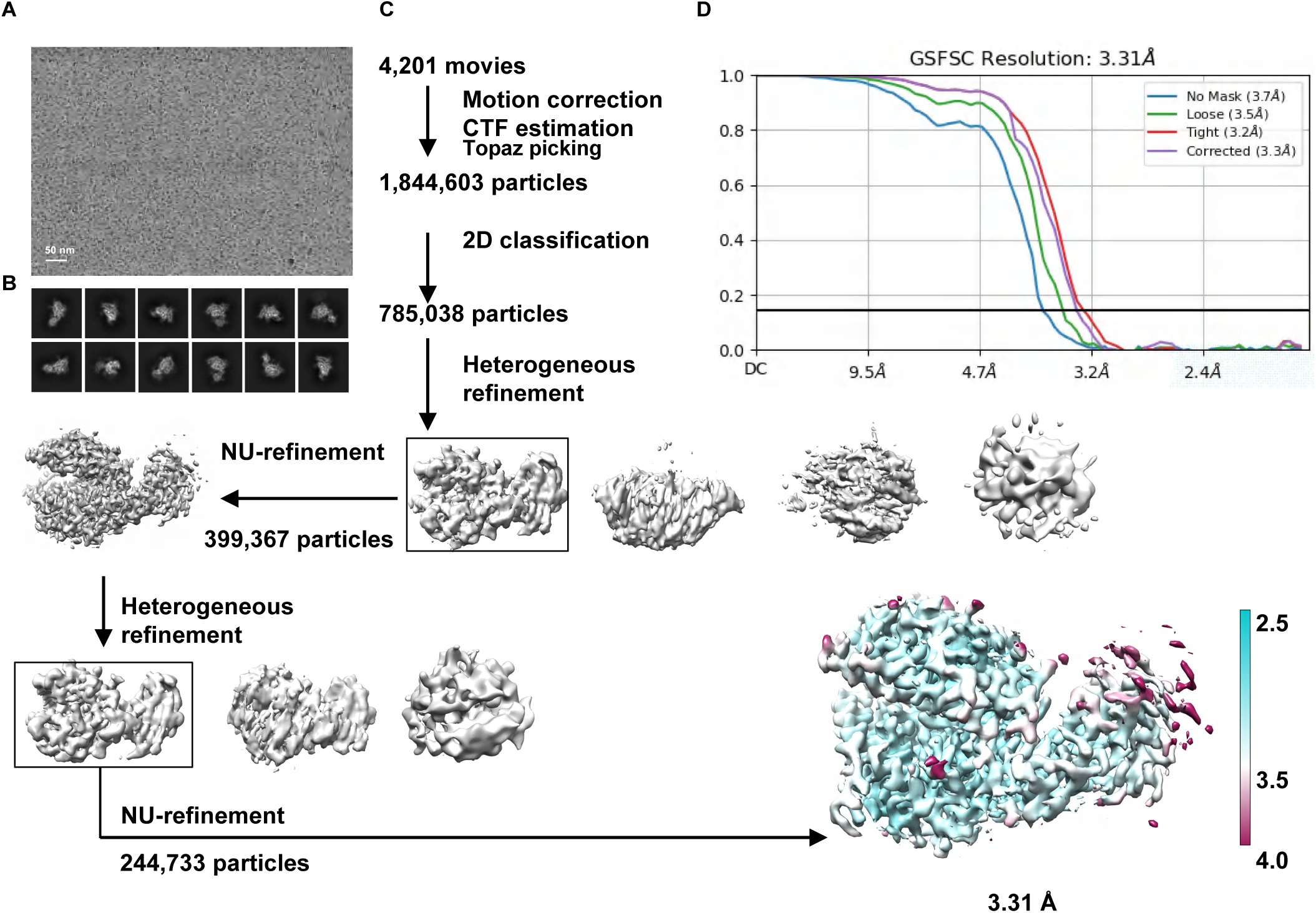
CryoEM data processing workflow for the P.nat ACE2.M2-bound MOW15-22 **RBD complex, related to** Figure 6. (**A-B**) **Representative electron micrograph and 2D class. averages of the P.nat ACE2.M2 bound MOW15-22 RBD complex embedded in vitreous ice. (C) Flowchart of cryo-EM data processing. (D) Fourier shell correlation (FSC) curve was calculated using two independent half maps, and resolution was estimated using the FSC=0.143 cutoff.**

**Figure S10.**
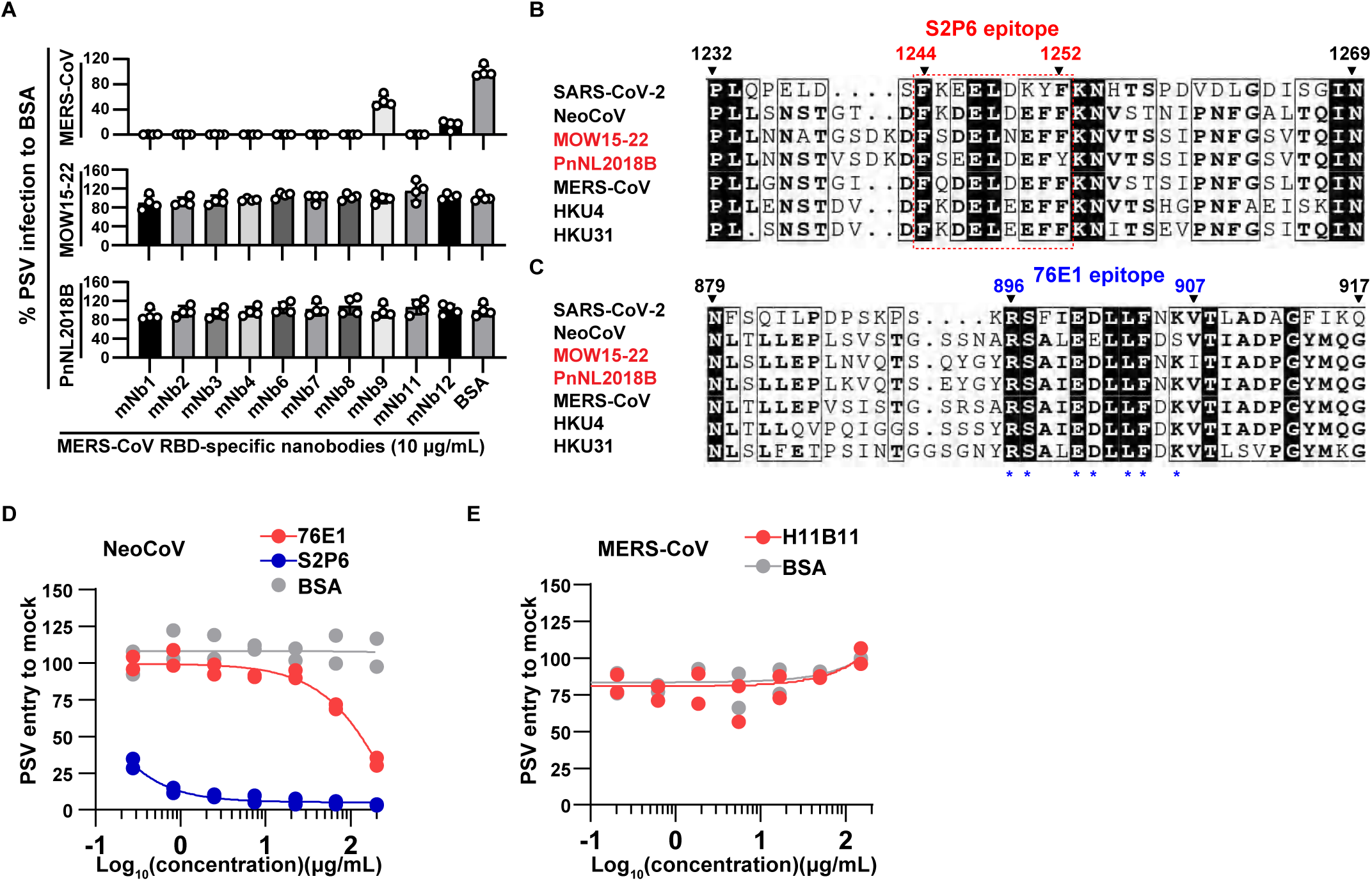
The neutralizing activity of monoclonal antibodies against PSV entry of indicated merbecoviruses, related to. Figure 6. (**A**) Neutralization efficiency of MERS-CoV RBD-directed nanobodies against MERS-CoV and MOW15-22 pseudotyped viruses. MEAN ± SD and unpaired two-tailed t-tests. n=3 biological replicates. (**B-C**) Sequence alignment displaying corresponding sequences of S2P6 (B) and 76E1 (C) epitopes of indicated coronaviruses. Red dashed box: S2P6 epitope. The MOW15-22 residue numbering is shown in B and C. (**D**) Dose-dependent inhibition of NeoCoV pseudovirus entry by antibodies targeting the stem helix (S2P6) or the S2’/fusion peptide (76E1) in HEK293T cells expressing P.pip ACE2. (**E**) The hACE2-specific antibody H11B11 does not inhibit MERS-CoV pseudovirus entry in Caco2-P.nat ACE2.3M cells. n=2 biological replicates for D and E.

**Table S1.**
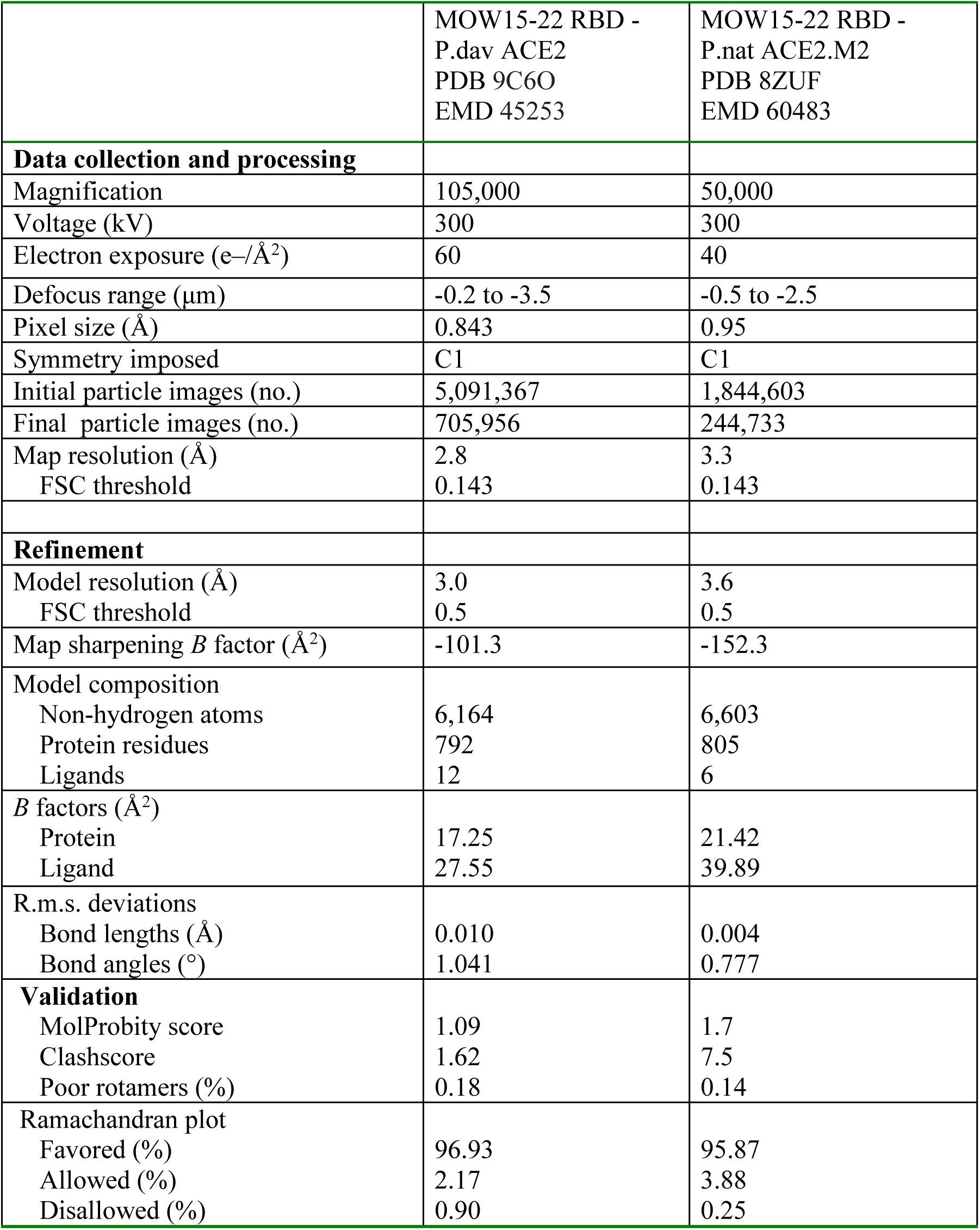
CryoEM data collection, processing, and model refinement statistics, related to. **Figure 5**.

**Table S2.** Conservation of receptor-interacting residues between MOW15-22 and PnNL2018B, related to. **Figure 5**.

| **MOW15-22 residue number** | **MOW15-22 residue identity** | **PnNL2018B residue identity** |
| --- | --- | --- |
| **505** | **G** | **G** |
| **506** | **C** | **C** |
| **507** | **V** | **V** |
| **508** | **D** | **G** |
| **509** | **K** | **Q** |
| **511** | **P** | **P** |
| **519** | **I** | **T** |
| **520** | **C** | **C** |
| **521** | **I** | **L** |
| **522** | **P** | **E** |
| **523** | **E** | **E** |
| **524** | **F** | **F** |
| **527** | **E** | **F** |
| **535** | **K** | **K** |
| **563** | **D** | **D** |
| **564** | **S** | **S** |
| **565** | **T** | **V** |
| **567** | **D** | **D** |
| **569** | **I** | **I** |
| **570** | **W** | **W** |
| **571** | **R** | **R** |

